# Two distinct mechanisms of small molecule inhibition of LpxA acyltransferase essential for lipopolysaccharide biosynthesis

**DOI:** 10.1101/850305

**Authors:** Wooseok Han, Xiaolei Ma, Carl J. Balibar, Christopher M. Baxter Rath, Bret Benton, Alun Bermingham, Fergal Casey, Barbara Chie-Leon, Min-Kyu Cho, Andreas O. Frank, Alexandra Frommlet, Chi-Min Ho, Patrick S. Lee, Min Li, Andreas Lingel, Sylvia Ma, Hanne Merritt, Elizabeth Ornelas, Gianfranco de Pascale, Ramadevi Prathapam, Katherine R. Prosen, Dita Rasper, Alexey Ruzin, William Sawyer, Jacob Shaul, Xiaoyu Shen, Steven Shia, Micah Steffek, Sharadha Subramanian, Jason Vo, Feng Wang, Charles Wartchow, Tsuyoshi Uehara

## Abstract

The lipopolysaccharide biosynthesis pathway is considered an attractive drug target against the rising threat of multidrug-resistant Gram-negative bacteria. Here, we report two novel small-molecule inhibitors (compounds **1** and **2**) of the acyltransferase LpxA, the first enzyme in the lipopolysaccharide biosynthesis pathway. We show genetically that the antibacterial activities of the compounds against efflux-deficient *Escherichia coli* are mediated by LpxA inhibition. Consistently, the compounds inhibited the LpxA enzymatic reaction in vitro. Intriguingly, using biochemical, biophysical, and structural characterization, we reveal two distinct mechanisms of LpxA inhibition; compound **1** is a substrate-competitive inhibitor targeting apo LpxA and compound **2** is an uncompetitive inhibitor targeting the LpxA-product complex. Compound **2** exhibited more favorable biological and physicochemical properties than compound **1**, and was optimized using structural information to achieve improved antibacterial activity against wild type *E. coli*. These results show that LpxA is a promising antibacterial target and imply the advantages of targeting enzyme-product complexes in drug discovery.

## INTRODUCTION

Bacterial antibiotic resistance is a growing crisis worldwide. In particular, untreatable and hard-to-treat infections by multidrug-resistant Gram-negative bacteria are a serious issue as they are becoming prevalent among patients in medical facilities.^1–2^ Multi-drug resistance mechanisms in Gram–negative pathogens consist mainly of antibiotic-modifying enzymes, active efflux, loss of outer membrane porins, target mutations, as well as combinations of these factors.^3^ Depending on their structure-activity relationships, novel antibacterial agents directed at new cellular targets could address such mechanisms of clinical resistance. However, target-based biochemical assays have only rarely provided leads with potent antibacterial activity against wild-type Gram-negative bacteria.^4–5^ This is because efflux and permeability barriers make many intracellular targets inaccessible to small molecule enzyme inhibitors. One strategy to address the limitation is to directly inhibit the lipopolysaccharide (LPS) biogenesis and transport pathways which enable these barriers.^6–16^

LPS, also known as endotoxin, is composed of a lipid A anchor that forms the outer leaflet of the outer membrane, a core oligosaccharide, and covalently-linked repeating polysaccharide O-antigen that extends out from the cell surface.^17^ Lipid A is produced by enzymes localized in the cytoplasm or the inner leaflet of the cytoplasmic membrane.^17^ Since lipid A is essential for growth and structural integrity of most Gram-negative bacteria, small molecule inhibitors of lipid A biosynthesis prevent growth or restore the activity of antibiotics with intracellular targets that could not otherwise be reached.^14–15, 18^ Furthermore, inhibition of lipid A biosynthesis can reduce the levels of endotoxin that are released during antibiotic treatment.^19^ Therefore, lipid A biosynthesis has been viewed as a promising target for the discovery of new antibacterials. The most advanced programs targeting lipid A are inhibition of LpxC, the second enzyme of the pathway.^14^ One of LpxC inhibitors reached clinical trials, however the development was hampered due to toxicity.^12^ Inhibition of other enzymes in the lipid A biosynthesis pathway is also appealing, but only a few small molecule inhibitors have been reported.^16, 20^

LpxA UDP-*N*-acetylglucosamine acyltransferase is the first enzyme in the lipid A biosynthesis pathway.^17^ This enzyme is conserved and essential in difficult-to-treat Gram-negative pathogens, with the exception of *Acinetobacter baumannii* which can grow without LPS under laboratory growth conditions.^21–22^ In *E. coli*, LpxA catalyzes the transfer of *R*-3-hydroxymyristate from acyl-carrier-protein to the 3-hydroxyl group of UDP-*N*-acetylglucosamine (UDP-GlcNAc), generating UDP-3-O-acyl-GlcNAc (Fig. 1a).^17^ Remarkably, the reaction catalyzed by *E. coli* LpxA is reversible with an unfavorable equilibrium constant (*K*_eq_ ≈ 0.01).^23–24^ LpxA contains an unusual, left-handed parallel β-helix fold and forms a soluble stable homotrimer constituting three active site pockets at the subunit interfaces.^25–27^ Crystal structures of *E. coli* LpxA in complex with its substrate UDP-GlcNAc^28^, product UDP-3-*O*-(*R*-3-hydroxymyristoyl)-GlcNAc^28^, and peptide inhibitors^29–33^ have been previously described, providing important insights into the catalytic mechanism and target validation. One of the peptide inhibitors is used as a tool molecule for development of assays to screen for compounds that bind LpxA.^34^ However, these peptide inhibitors cannot be chemical starting points for development of antibacterial agents targeting cytoplasmic LpxA due to the lack of the delivery system. Although small molecule inhibitors against *Pseudomonas aeruginosa* LpxA and LpxD were reported recently^16^, no LpxA inhibitor having antibacterial activity has been publicly disclosed.

**Figure 1.**
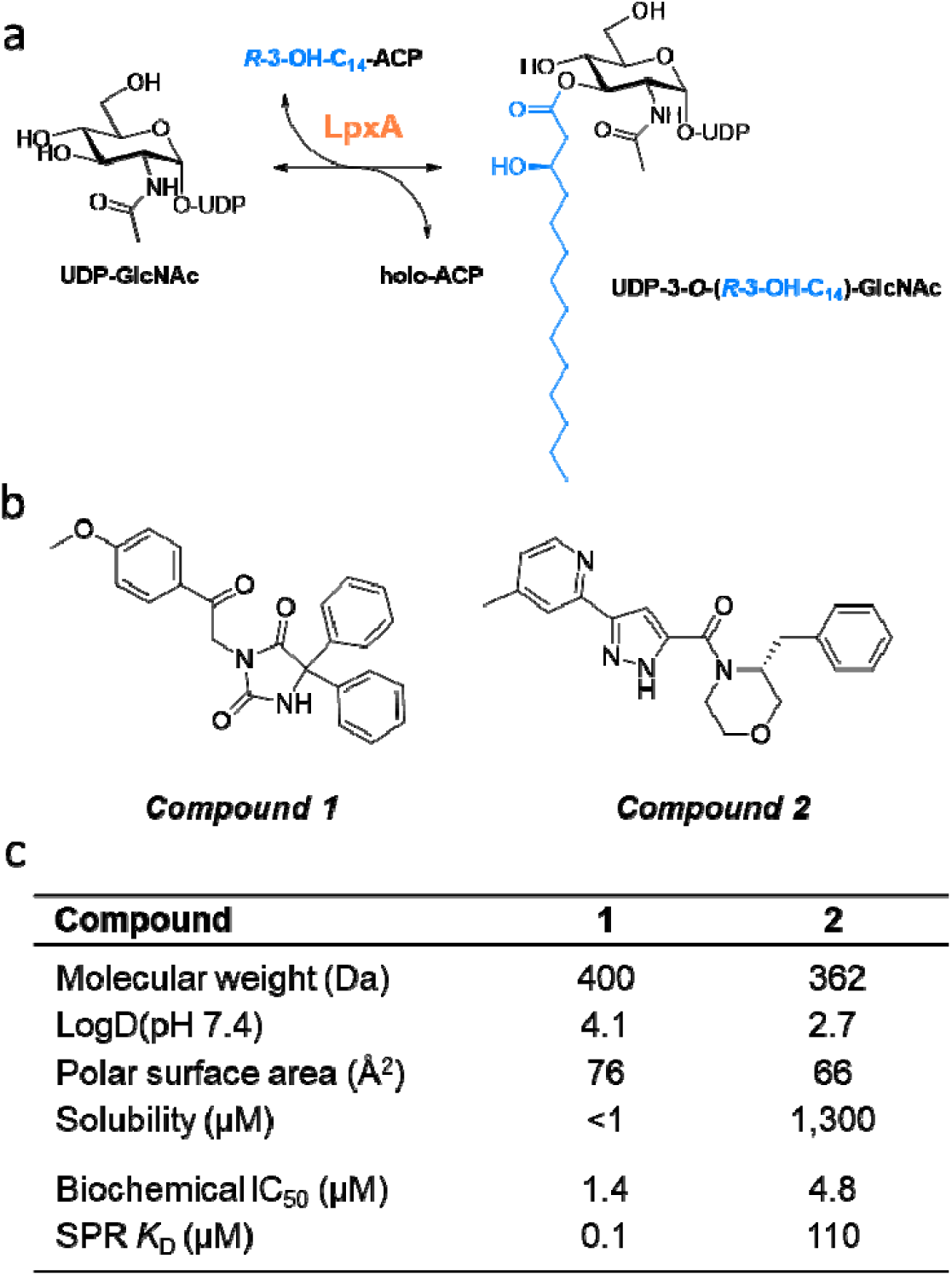
*E. coli* LpxA reaction and the chemical structures and properties and biological activities of compounds 1 and 2. **(a)** *E. coli* LpxA reaction. LpxA utilizes *R*-3-hydroxy-myristoyl-acyl-carrier protein (*R*-3-OH-C_14_-ACP) to transfer the fatty acyl chain to UDP-GlcNAc, generating UDP-3-*O*-(*R*-3-OH-C_14_)-GlcNAc. **(b)** Chemical structures of compounds **1** and **2. (c)** Chemical properties and in vitro activities of the compounds.

Here we present the small molecule inhibitors of *E. coli* LpxA, compounds **1** and **2** (Fig. 1b). The compounds are among the hits from cell-based screenings of Novartis compound collections for bacterial growth inhibitors and show antibacterial activity against efflux-deficient *E. coli* (Δ*tolC*) with no measurable activity of eukaryotic cell cytotoxicity or hemolysis at 100 μM. Genetic target identification and in vitro inhibitory activity in the LpxA-specific biochemical assay indicated that both compounds were LpxA inhibitors with antibacterial activity mediated by the target inhibition. Intriguingly, we found that the compounds showed distinct mechanisms of inhibition. Using biochemical and biophysical characterizations and X-ray crystallography, we demonstrate that compound **1** is an inhibitor of apo LpxA and that compound **2** inhibits LpxA only in the presence of the product UDP-3-*O*-(*R*-3-hydroxymyristoyl)-GlcNAc. Compound **2** was prioritized for follow up and optimized to achieve a minimal inhibitory concentration (MIC) of 16 µg/mL against wild type *E. coli*. Our findings establish that the LpxA-product complex is a promising target for antibiotic discovery, and provide advantages to design product-dependent inhibitors for target-based drug discovery.

## RESULTS

### Compound 1 is an apo LpxA inhibitor

To identify the cellular target of the antibacterial active compound **1**, we performed mutant selection of *E. coli* Δ*tolC* cells and yielded a mutant that had a FabZ A146D substitution. Mutations in *fabZ* are known to suppress the growth defect of *lpxA* and *lpxC* mutants and also reduce the activity of LpxC inhibitors^35–37^ and an LpxD inhibitor^20^. Therefore, we postulated that compound **1** inhibited one or more of the Lpx enzymes rather than FabZ. Consistent with this, three *fabZ* mutants that encoded different FabZ substitutions were less susceptible to compound **1** and the LpxC inhibitor CHIR-090 (Table 1). We then examined whether overexpression of a selection of essential lipid A biosynthetic enzymes altered susceptibility to compound **1**. Susceptibility was reduced only by LpxA overexpression (Table 1), suggesting that the antibacterial activity of compound **1** was mediated by inhibition of LpxA. To show whether the compound directly inhibited LpxA, we tested compound **1** in an in vitro biochemical assay using a solid-phase-extraction mass-spectrometry (SPE-MS) based read out for the LpxA product. Compound **1** showed a half-maximal inhibitory concentration (IC_50_) of 1.4 μM, indicating that compound **1** is an LpxA inhibitor (Fig. 1c).

**Table 1.** MICs of the LpxA inhibitors and the LpxC inhibitor CHIR-090.

| Strain | Description | Compound 1 |  | Compound 2 |  | CHIR-090 |  |
| --- | --- | --- | --- | --- | --- | --- | --- |
|  |  | MIC | Fold shift | MIC | Fold shift | MIC | Fold shift |
| ATCC 25922 | *E. coli* clinical isolates | >16 | - | >128 | - | ND | - |
| MG1655 | *E. coli* K-12 | >16 | - | >128 | - | ND | - |
| VECO2526 | *E. coli* $\Delta tolC$ | 0.5 | 1 | 2 | 1 | 0.015 | 1 |
| TUP0115* | $\Delta tolC$ pLacZα↑ | 0.5 | 1 | 2 | 1 | 0.008 | 0.5 |
| TUP0116* | $\Delta tolC$ pLpxA↑ | **>16** | **>32** | **>64** | **>32** | **0.06** | **4** |
| TUP0118* | $\Delta tolC$ pLpxC↑ | 0.5 | 1 | 2 | 1 | **0.125** | **8** |
| TUP0119* | $\Delta tolC$ pLpxD↑ | 0.5 | 1 | 2 | 1 | 0.015 | 1 |
| ARA0016 | $\Delta tolC$ FabZ<sub>A78V</sub> | **>16** | **>32** | 1 | 0.5 | **0.25** | **16** |
| TUP0074 | $\Delta tolC$ FabZ<sub>P56L</sub> | **>16** | **>32** | 2 | 1 | **0.25** | **16** |
| TUP0042 | $\Delta tolC$ FabZ<sub>C145Y</sub> | **>16** | **>32** | 2 | 1 | **0.25** | **16** |
| CDY0154 | *E. coli* $\Delta 9efflux$ | 0.5 | 1 | 2 | 1 | 0.075 | 1 |
| TUP0093 | $\Delta 9efflux$ LpxA<sub>Q73L</sub> | 1 | 2 | **64** | **32** | 0.075 | 1 |
| TUP0092 | $\Delta 9efflux$ AccB<sub>E128K</sub> | 0.5 | 1 | **32** | **16** | 0.075 | 1 |
| TUP0097 | $\Delta 9efflux$ InfA<sub>R66H</sub> | 1 | 2 | **32** | **16** | 0.075 | 1 |

The values are modes of MIC values measured in at least two independent experiments. The MIC with asterisks was determined in the presence of 100 μM IPTG to induce the gene expression (up-arrow). TUP0115 carries a control vector expressing LacZ $\alpha$ . The unit of MIC values is μg/mL. Numbers in parentheses are fold shifts in MICs relative to those against the corresponding parent strains. The MIC fold shifts with $\geq 4$ are shown in bold. Although compound **1** was tested at the maximum concentration of 128 μg/mL in the MIC assay, precipitation was observed at 32, 64, and 128 μg/mL. Due to the precipitation, >16 μg/mL is reported when no inhibition of growth was observed at 16 μg/mL and higher concentrations.

To determine the mechanism of inhibition of compound **1**, we tested the compound in an Surface plasmon resonance (SPR) binding assay to measure binding affinity for apo LpxA. We found that compound **1** bound surface-immobilized LpxA with a *K*_D_ of 0.1 µM (Fig. 2a), suggesting that this compound inhibits LpxA by binding the apo form. Enzymatic characterization was also performed using titration of each of the two LpxA substrates (acyl-ACP and UDP-GlcNAc). Compound **1** showed inhibition competitive with acyl-ACP and non-competitive to UDP-GlcNAc (Fig. 3), which is consistent with apo LpxA being its target. We further obtained the LpxA X-ray co-structure with compound **1** at 2.1 Å by soaking (Fig. 4a, S5). Compound **1** clearly bound to apo LpxA (three inhibitors per trimer) by occupying a large portion of the acyl chain binding pocket. The hydantoin motif of compound **1** engaged two adjacent LpxA subunits by H-bonding interactions to Q161 of one subunit and G155 of the other subunit. The binding site for the compound also included a smaller hydrophobic pocket lined by M118, I134, A136, and I152. Interestingly, G155 has previously been implicated in pantetheine binding to *Leptospira interrogans* LpxA^38^. Collectively, these results reveal that compound **1** inhibits LpxA function by directly targeting the apo enzyme.

**Figure 2.**
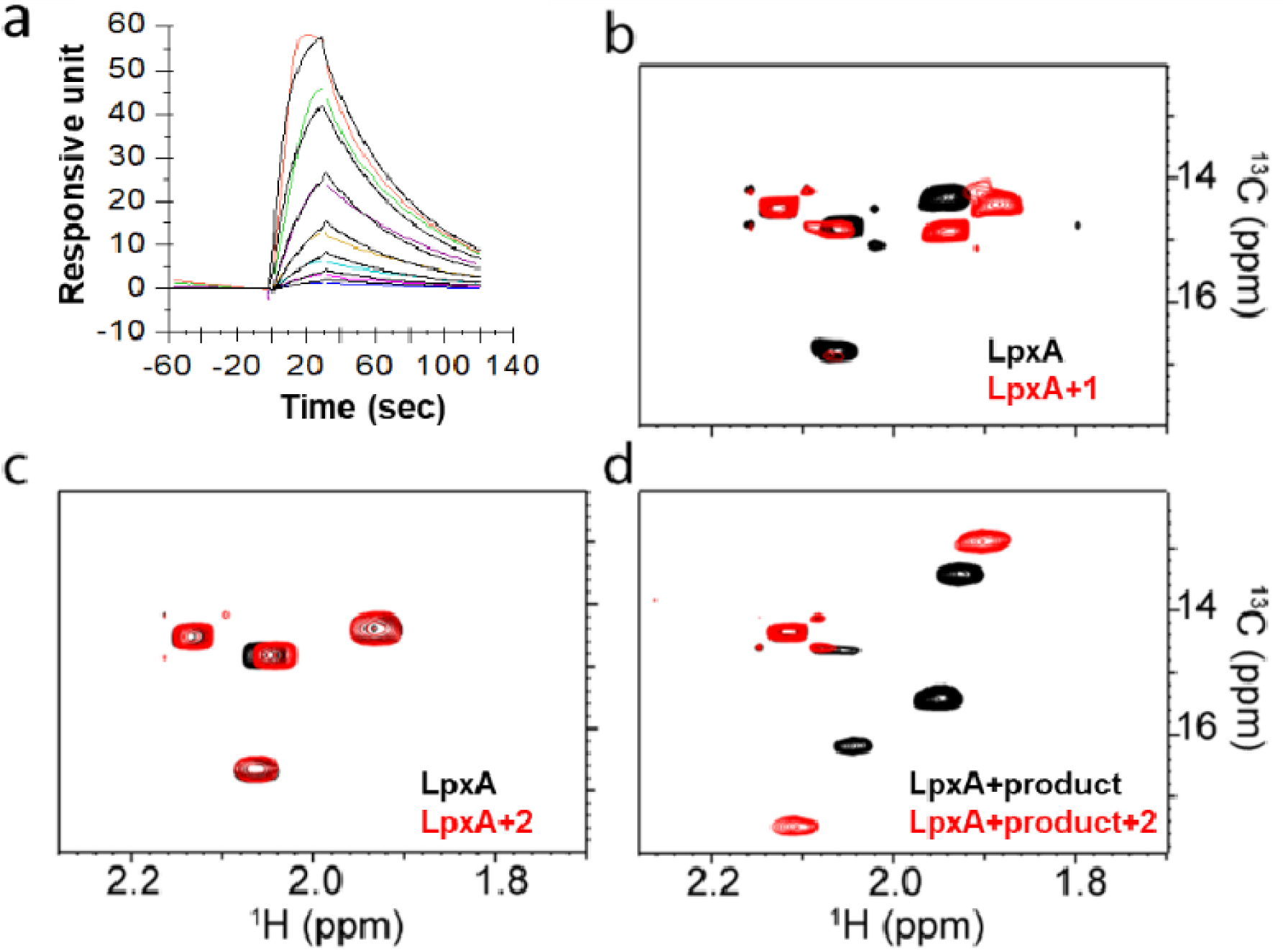
Biophysical characterization of compounds 1 and 2. **(a)** SPR sensorgram. Avi-tagged LpxA was immobilized to the SPR chip surface. The responsive units for compound **1** (6 point concentrations, 2× dilutions, top 100 µM) plotted are shown over time. The kinetic K_D_ of compound **1** determined with a 1:1 binding was 0.1 µM.SPR sensorgram for compound **2** is shown in Fig. S3. **(b, c, d)** Protein-observed 2D ^1^H-^13^C HMQC (heteronuclear multiple quantum correlation) spectra of the labeled amino acid residues (MILVAT) of LpxA. Shown are the peaks representing the methyl group of the LpxA methionine side chain in the absence (black) or presence (red) of compound **1** (b) or compound **2** (c). LpxA peaks with the product UDP-3-*O*-(*R*-3-hydroxymyristoyl)-GlcNAc are shown in the absence (black) or presence (red) of compound **2** (d). Peak shifts represent ligand binding to LpxA. The concentrations in samples were 80 µM isotope-labeled LpxA, 100 µM compound, and 150 µM LpxA product. The entire methyl regions of the spectra are shown in Fig. S4.

**Figure 3.**
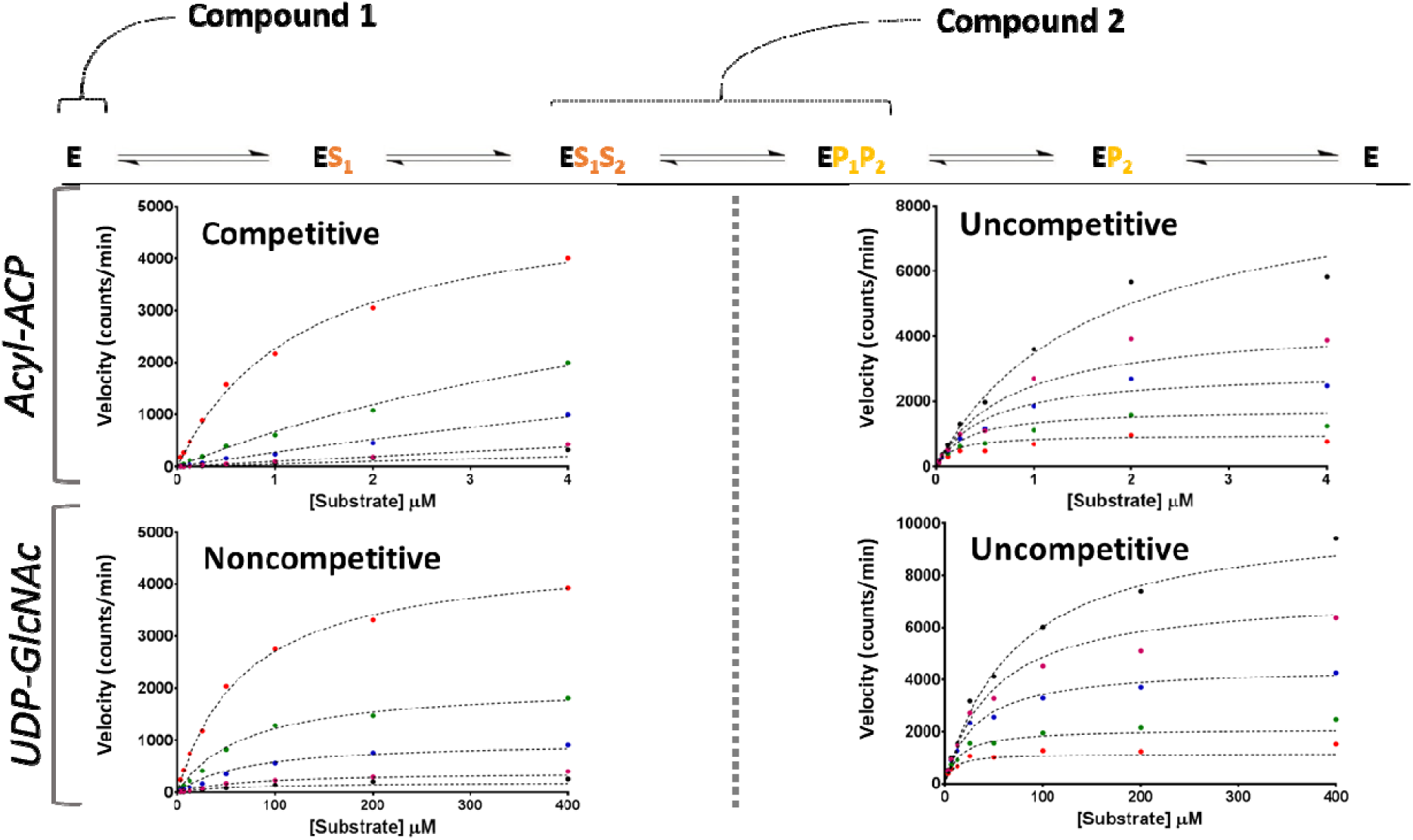
Biochemical characterization of compounds 1 and 2. The scheme of the enzymatic steps in the LpxA reaction is shown on the top. (Left) compound **1** showed competitive inhibition to acyl-ACP and non-competitive to UDP-GlcNAc. (Right) compound **2** showed uncompetitive inhibition to both substrates. The inhibitor concentrations used were 0, 0.48, 1.45, 4.35, and 8.7 µM for compound **1**, and 0, 1.3, 4.1, 12.3, and 25 µM for compound **2**. Data are mean of two assays. The signals (counts) of the product were measured using SPE-MS.

**Figure 4.**
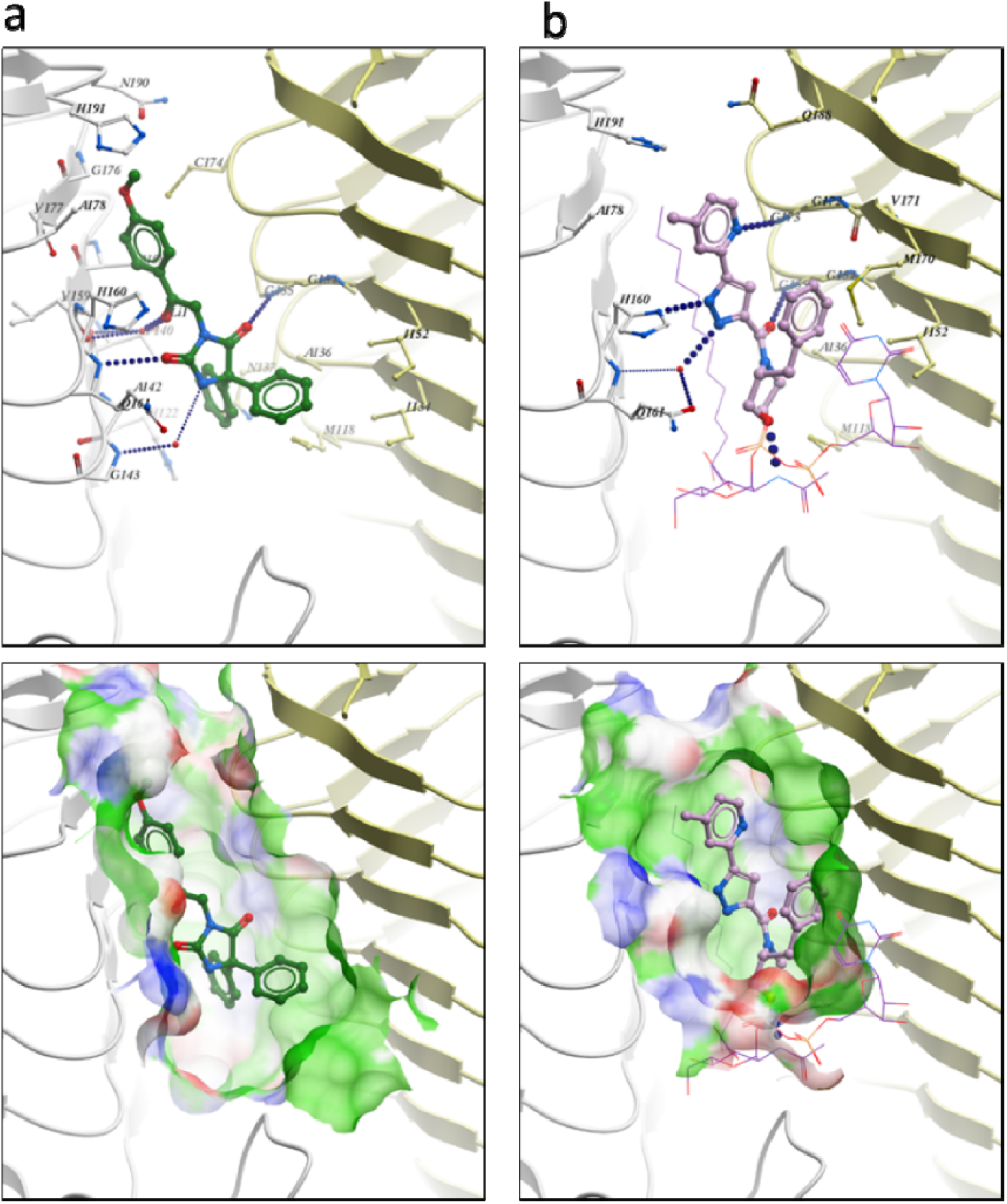
Co-structures of LpxA in complex with compound 1 and 2. The X-ray crystal structures in complex with compound **1** (a) and with the product and compound **2** (b) are shown. The asymmetric unit contains one LpxA monomer in complex with one LpxA product (UDP-3-*O*-(*R*-3-hydroxymyristate)-GlcNAc) and one compound, and the biologically functional trimer lies on the crystallographic 3-fold axis. LpxA backbone is shown with carbon colored white for subunit 1, yellow for subunit 2. LpxA residues near ligands bound are shown as cartoon ball and sticks representation, oxygen colored red and nitrogen colored blue. Compounds are shown as ball and sticks with carbon colored green (compound **1**) and purple (compound **2**). LpxA product (b) is shown as sticks with carbon colored purple. (Bottom) The binding pockets for the compounds are shown with the representation of hydrophobic (green), positive-charged (red), and negative-charged (blue) surface.

### Compound 2 also targets LpxA in *E. coli*

To identify the cellular target of compound **2**, we utilized a gene overexpression library in the *E. coli* strain lacking nine efflux pumps (CDY0154). The library was constructed using the mobile plasmid collection^39^ where every *E. coli* gene is expressed from an inducible promoter of a low copy plasmid. Selection by compound **2** yielded only the *lpxA* clone in 40 separately-isolated colonies, suggesting that compound **2** was also an LpxA inhibitor. This was further confirmed with reduced susceptibility by overproduction of LpxA expressed from a different plasmid backbone, but not by LpxC or LpxD (Table 1). We then tested the impact of *fabZ* mutations on compound susceptibility. Unexpectedly, the *fabZ* mutant strains were equally susceptible to compound **2** in contrast to their reduced susceptibility to compound **1** and the LpxC inhibitor CHIR-090 (Table 1).

To further understand the cellular target of compound **2**, we isolated mutants with reduced susceptibility by passaging CDY0154 cells in the presence of compound **14**, an analog of compound **2** (Fig. S2). Among a total of 18 colonies isolated by mutant selection, three mutants were used for genome sequencing and susceptibility tests. Mutant TUP0093 had an LpxA Q73L substitution and was 32-fold less susceptible to compound **2** (Table 1). The other two mutants were 16-fold less susceptible and had mutations in *accB* or *infA* (Table 1). InfA is an initiation factor for translation^40^ and AccB is a component of the Acetyl-CoA Carboxylase (Acc) complex that catalyzes the first reaction of fatty acid biosynthesis^41^. Neither the *accB* nor the *infA* mutation affected the MICs of compound **1** or CHIR-090 (Table 1), which together with the susceptibility of the *fabZ* mutants suggested a specific mechanism to reduce susceptibility to compound **2**. Since *accB* and *infA* are also essential for growth^42^, we cannot exclude the possibility that compound **2** inhibits one or both protein functions. However, compound **2** showed direct inhibition of the in vitro LpxA reaction with an IC_50_ of 4.8 µM (Fig. 1c), indicating that compound **2** is an LpxA inhibitor. The distinct resistant profiles suggest that the two inhibitors, compounds **1** and **2**, differed in their mode of LpxA inhibition.

### Compound 2 is a product-dependent LpxA inhibitor

Despite the biochemical IC_50_ of 4.8 µM, compound **2** bound weakly to apo LpxA (SPR *K*_D_ = 110 µM, Fig. 1c, S3). This discrepancy between the biophysical and biochemical potency of compound **2** prompted us to further investigate its underlying mechanism of action. Enzyme kinetics studies showed that compound **2** was uncompetitive with both LpxA substrates (Fig. 3), consistent with the weak binding to apo LpxA. This result also indicated dependence on the presence of either a substrate or a product for inhibition by compound **2**. Based on the unfavorable forward reaction of LpxA^23^ and the stable enzyme/product complex demonstrated by the crystal structure with UDP-3-*O*-(*R*-3-hydroxymyristoyl)-GlcNAc^28^, we postulated that compound **2** might bind the LpxA-product complex. To examine the binding mechanism, we developed a two-dimensional protein-observed HMQC NMR assay^43–44^ for *E. coli* LpxA, using selective incorporation of ^1^H,^13^C isotope labels at all methyl positions of CH_3_-containing amino acids (Met, Ile, Leu, Val, Ala, and Thr, but not Ile Cγ). As expected, robust chemical shift perturbations (CSP) were observed when isotope-labeled LpxA was incubated with compound **1** (Fig. 2b). In contrast, few small CSPs of LpxA resonances were observed upon addition of 100 µM of compound **2**, consistent with weak binding to apo-LpxA (Fig. 2c). As we hypothesized, drastic CSPs and signal broadening were observed for product-bound LpxA in the presence of compound **2** (Fig. 2d, S4). These results indicate that compound **2** inhibits the LpxA reaction in a product-dependent manner.

### LpxA crystal structure in complex with product UDP-3-*O*-(*R*-3-hydroxymyristoyl)-GlcNAc and compound 2

To further validate and characterize the LpxA/product complex as the molecular target of compound **2**, unliganded LpxA crystals were soaked with 2 mM compound **2** and 40 mM UDP-3-*O*-(*R*-3-hydroxymyristoyl)-GlcNAc. The crystals diffracted to 1.8 Å resolution with excellent geometry as assessed with Mol-Probity^45^ (Table S3). Compared to the LpxA structure in complex with its product^28^, additional electron density that fitted compound **2** was clearly observed and belonged to the *R*-enantiomer (Fig. S5), representing the biologically active form of compound **2**. Compound **2** occupied the pantetheine site and formed extensive interactions with both the LpxA protein and LpxA product UDP-3-*O*-acyl-GlcNAc (Fig. 4b). Intriguingly, five of six heteroatoms of compound **2** were involved in direct or water mediated H-bonds. Specifically, the pyridine and pyrazole moieties formed direct H-bonds with the backbone carbonyl of G173 and H160 of LpxA, respectively. The pyrazole formed a water-mediated interaction with Q161. The morpholine moiety was directly associated with the nitrogen atom at the C2 position of the GlcNAc moiety of the product. The benzyl group mediated hydrophobic interactions with LpxA M170 and I152. This structure reveals why compound **2** shows a low affinity for apo LpxA in SPR and NMR, and is consistent with product-dependent inhibition.

### Structure-based compound design improved the antibacterial activity

Although the LpxA inhibitor compounds **1** and **2** exhibited good activity against the efflux-deficient *E. coli* strains, both did not inhibit growth of the *E. coli* ATCC 25922 clinical isolate even at 128 µg/mL (Table 1). To improve cellular activity against wild type *E. coli*, we initiated structure-based optimization of compound **2** with favorable properties for Gram-negative antibacterial agents that must pass through bacterial outer and inner membranes to reach their targets^46^; that is, compound **2** had higher solubility, lower logD (pH 7.4), and lower molecular weight than compound **1** (Fig. 1c). Furthermore, the binding pocket of the LpxA-product complex was much smaller and more polar than that of the apo enzyme (Fig. 4). These attributes made compound **2** the more progressible chemical starting point for optimization for potency against wild type *E. coli*.

We designed and synthesized analogs of compound **2** using structural information and in consideration of the physicochemical properties preferred for antibiotics that are active against Gram-negative bacteria.^46–47^. In the X-ray co-crystal structure of compound **2** and the LpxA-product complex, the electron density of the benzyl moiety was weaker than that of the other parts of molecule. This indicated that the benzyl group was partly flexible. Therefore, we expected that optimization in this region could increase the inhibitory potency while maintaining or improving physicochemical properties. Removal or replacement of the benzyl with alkyl groups and heterocyclic rings, such as pyridines and pyrazoles (compounds **3, 4**, and **5**) resulted in a significant loss of potency in biochemical and MIC assays (Fig. 5), suggesting that the benzyl moiety is important for maintaining the antibacterial effect. With respect to substitutions to the benzyl ring, ortho-substituents favorably influenced antibacterial activity in comparison with meta- or para-substituents. In particular, the addition of a fluoro group at the ortho-position of the benzyl group (compound **6**) led to 4-fold improvement in MIC against the efflux-deficient *E. coli* Δ*tolC*. Likewise, compound **7** having ortho-chloro substituent and compound **8** having ortho-methoxy substituent increased an antibacterial effect on *E. coli* Δ*tolC*. To evaluate the effect of the ortho-substitution on compound binding, we obtained LpxA co-crystal structures with compounds **6** and **8** in the presence of UDP-3-*O*-(*R*-3-hydroxymyristoyl)-GlcNAc. The structures showed that the ortho-substituents in compounds **6** and **8** were located in different regions: the fluoro group in compound **6** faced toward the protein and the methoxy group in compound **7** was exposed to solvents (Fig. 6a, Supplementary Fig. 5). In both cases, the benzyl ring was shifted toward the protein by pulling or pushing the ring, respectively. In addition, the ortho-substituted benzyl groups were much better defined in the electron density map compared to the benzyl group of compound **2** (Supplementary Fig. 5). Based on this structural information, we prepared compounds **9** and **11** (Fig. 5) that had substituents at the two different ortho-positions. Compound **11** having chloro and methoxy substituents at the ortho-positions showed better biochemical IC_50_ than compound **9** having fluoro and methoxy substituents, suggesting a better ortho-substitution of chloro than fluoro for ligand binding. Compound **11** exhibited slightly better antibacterial activity against a wild type clinical isolate, *E. coli* ATCC 25922 (MIC = 32 µg/mL) than compound **9**. Furthermore, polar functional group at the ortho position of the methyl in the pyridine ring was anticipated to make a hydrogen bond interaction with Q188. Compound **13** having an amino group in the pyridine ring showed an MIC of 16 µg/mL against wild type *E. coli*. Using the X-ray crystallography, we confirmed a hydrogen bond interaction of the amino group of compound **13** made with Q188 of one subunit of LpxA and found additional interaction with His191 of the other subunit (Fig. 6b). The antibacterial activities of these analogs were very likely to be mediated by LpxA inhibition because of the reduction of cell susceptibility by the LpxA Q73L variant (Fig. 5) and the less activities shown by the enantiomers of compound **2** and **9** (compound **12** and **10**, respectively). Consistently, we found a moderate linear correlation between the biochemical inhibition and the antibacterial activity against *E. coli* Δ*tolC* for analogs of compound **2** (Fig. 6c). Taken together, the introduction of three different functional groups, chloro, methoxy, and amino groups resulted in an 8-fold improvement in both biochemical inhibition and antibacterial activity, leading to an MIC of 16 µg/mL against wild type *E. coli*.

**Figure 5.**
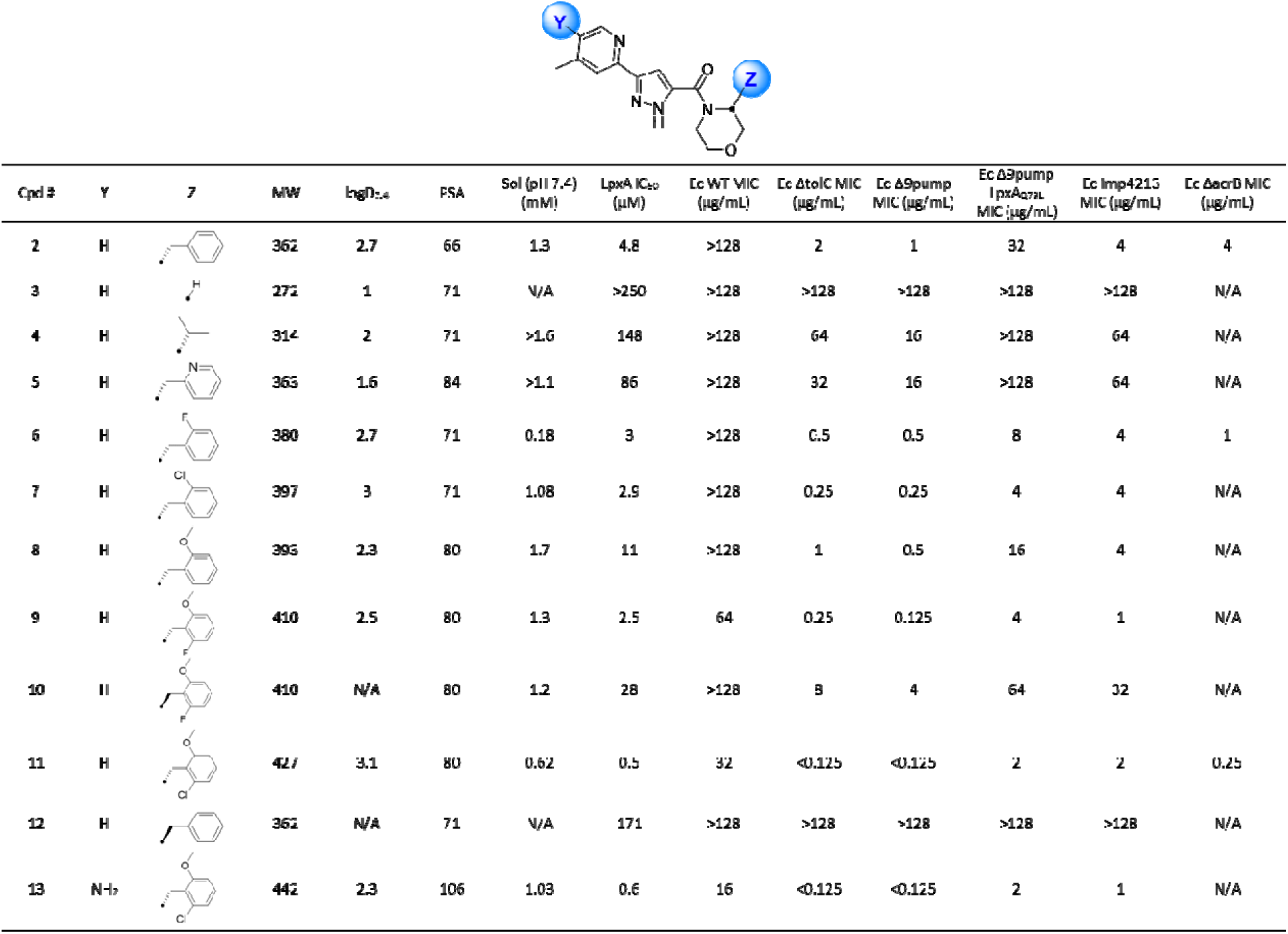
Physicochemical properties and biological activities of compound 2 and the analogs. MW, molecular weight; PSA, polar surface area; Sol, solubility; Ec, *E. coli*; ND, not determined; Ec WT, ATCC 25922; Ec Δ*tolC*, JW5503-1; Ec Δ*acrB*, JW0451. Compound **10** (compound **9** enantiomer) was 94% pure after chiral separation. As it contained a small amount of compound **9**, it showed weak biological activities. Compound **12** (compound **2** enantiomer) was >99% pure.

**Figure 6.**
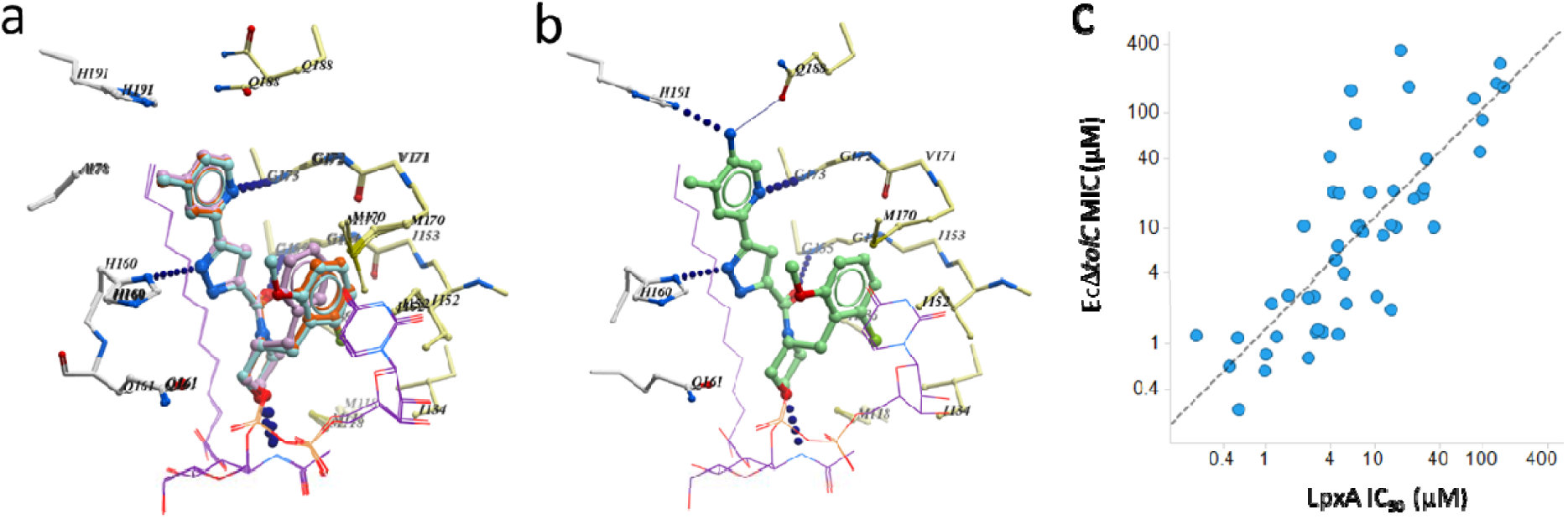
Structure-guided optimization of compound 2. (a) An overlay of the LpxA crystal structures in complex with compounds **2** (purple), **6** (red), and **8** (aqua) and with the product. The ortho fluoro group (compound **6**) and the ortho methoxy group (compound **8**) were observed in opposite orientations in the structures. (b) The X-ray crystal structures in complex with the product and compound **13**. (c) Correlation of in vitro enzyme inhibition (LpxA IC_50_) and MIC against *E. coli* Δ*tolC* for LpxA inhibitors (n = 49, blue circle). Compounds that showed >250 uM IC_50_ or >128 ug/mL EcΔ*tolC* MIC were not plotted. A simple linear regression is shown as dashed line (*R*^*2*^ = 0.635).

## DISCUSSION

Here we report novel LpxA small molecule inhibitors. Notably, the inhibitors had two distinct mechanisms of action: the substrate-competitive inhibitor compound **1** and the product-dependent inhibitor compound **2**. While substrate-competitive inhibitors are commonly identified in target-based screenings, product-dependent inhibitors are more unusual. An example is triclosan, the antibiotic that inhibits FabI only in the presence of the product NAD^+^ and forms a tight ternary complex^48^. Overall, the product-dependent inhibitor compound **2** showed more favorable biological and chemical properties than compound **1**.

Resistance profiling was one way to distinguish between the two LpxA inhibition mechanisms. Mutations in *fabZ* decreased susceptibility to compound **1** as well as to inhibitors of LpxC and LpxD. This is probably because altered FabZ proteins are less active, resulting in an increased cellular level of *R*-3-hydroxymyristoyl-ACP which competes with compound **1** for binding to LpxA. This is analogous to the previously-proposed mechanism of resistance by *fabZ* mutations to the substrate-competitive LpxC inhibitor CHIR-090.^35^ In contrast, *fabZ* mutations did not affect the potency of compound **2**, supporting its substrate-uncompetitive mechanism. Therefore, the increased metabolic flux to LPS biosynthesis that occurs in *fabZ* mutants should increase the IC_50_ of substrate-competitive inhibitors of LpxA, LpxC, and LpxD, but not influence the activity of the substrate-uncompetitive product-dependent inhibition of LpxA. LpxC inhibitors are in development for treatment of Gram-negative pathogens^8, 14, 49^, and if an LpxC inhibitor is approved and used in the clinic, pathogens would be expected to acquire *fabZ* mutations over time. These mutants would become cross-resistant to an apo LpxA inhibitor, but still susceptible to a product-dependent LpxA inhibitor.

The *lpxA* missense mutation that reduced compound **2** activity caused the substitution of Gln73 to Leu. The side chain of Q73 is located in the active site pocket and forms a direct hydrogen bond with the 3□-hydroxyl group of the product.^28^ It was therefore unexpected that Q73 was not essential for LpxA activity and bacterial viability. There was no direct interaction between Q73 and compound **2** in the ternary complex structure, suggesting that Q73L may affect the configuration of product binding to LpxA, thereby reducing the affinity of compound **2** to the LpxA-product complex. The LpxA Q73L variant did not affect compound **1** activity, consistent with no direct interaction being present between compound **1** and Q73 in the LpxA-compound **1** co-structure. The difference in susceptibility profiles for the two LpxA inhibitors make the LpxA_Q73L_ mutant a useful tool for validating cellular on-target activity of chemical modifications around the compound **2** scaffold.

Using structure-based inhibitor design, we could improve the antibacterial and biochemical activities of compound **2** by 8 fold. The best compound (compound **13**) in the series showed sub-micro molar IC_50_ and an MIC of 16 µg/mL against *E. coli* ATCC 25933, a clinical isolate. It is possible that the affinity for LpxA in the product-dependent inhibition mode is saturated or that it is limited by the affinity of the LpxA product. Product-dependent inhibition of FabI by triclosan is very potent (IC_50_ in the nanomolar range), while the affinity of NAD^+^ for FabI is in the low millimolar range.^48, 50^ In the case of triclosan, therefore, the potency of product-dependent inhibition does not depend on product affinity. It remains unclear whether the affinity of compound **2** is dependent on product affinity to LpxA. In the LpxA ternary complex, compound **2** interacted with both the product and residues in the LpxA active site. Thus, further optimization of product-dependent inhibitors for binding to LpxA protein residues is expected to lead to significant increases in affinity. In addition to affinity enhancement, compound **2** series need further improvement in the MICs against clinical isolates of *E. coli* (and other Gram-negative pathogens) in order to move the LpxA inhibitor forward on lead optimization. In general, understanding the chemical properties that facilitate compound accumulation in Gram-negative bacterial cells by overcoming outer membrane permeability and efflux is important for antibacterial discovery^4, 51–53^. However, universal guidelines that can be used to optimize compounds for better bacterial cell accumulation are not yet established because molecular descriptors for compound accumulation vary for each chemical scaffold^53^. Understanding of cell entry for this chemical series as well as insights into increasing binding-affinity to the LpxA-product complex are necessary to further increase potency and accumulation in *E. coli*.

In conclusion, the identification of the inhibitors with the two distinct mechanisms of action and the improvement of the potency of the product-dependent inhibitor exhibit that LpxA is a promising antibacterial target against multi-drug resistant Gram-negative bacteria. This work also shows that LpxA overexpression is a useful method to identify compounds that target LpxA with various mechanisms of action in the cellular context. This genetic method is based on antibacterial activity of the compound and cannot be applied to identify inhibitors that are inactive or weakly active against efflux-deficient *E. coli*, nor those unaffected by LpxA overexpression (these compounds could potentially be identified by biochemical or biophysical assays, as will be reported elsewhere). A distinct success of the genetic, biochemical, biophysical, and structural work was the identification of a product-dependent inhibitor. The chemical and biological advantages of product-dependent inhibition can be applied to other target enzymes, especially those having relatively large active site pockets. The presence of a product or substrate in the active site(s) limits the size of the binding pocket, so that inhibitors can be more easily designed to fit the chemical property space required for the indication of interest.

## METHODS

Full methods are available in Supporting Information. Bacterial strains and plasmids used in this study are listed in Table S1. MICs were determined by the broth dilution method. Genetic target identification was performed using materials and methods including spontaneous resistant mutant isolation, the mobile plasmid collection, genome sequencing, PCR, and DNA sequencing. LpxA-His6, biotinylated Avi-LpxA, and holo-ACP were overexpressed in *E. coli* and purified by chromatographic steps with appropriate enzymatic treatments. Acyl-ACP was prepared by enzymatic acylation of holo-ACP and purification by chromatography. The in vitro LpxA biochemical reaction was performed by incubation of LpxA-His6 and the substrates for 15 min at room temperature, followed by SPE-MS detection of the generated LpxA product UDP-3-*O*-(*R*-3-hydroxymyristate)-GlcNAc using Agilent RapidFire 365 coupled to Sciex 5500 triple quadrupole MS. The LpxA SPR binding assay was performed using Biacore T200. Protein-observed NMR using isotope-enriched LpxA-His6 protein was performed as described previously.^43^ The X-ray crystal structures of the LpxA-ligand complexes were obtained by crystal soaking and solved by molecular replacement. Compound **1** and 5-(4-methylpyridin-2-yl)-1H-pyrazole-3-carboxylic acid for compound **2** were prepared using literature procedures.^54–55^ Morpholine intermediates were purchased or prepared using two methods. Compound **2** and the analogs were prepared by amide coupling of 5-(4-methylpyridin-2-yl)-1H-pyrazole-3-carboxylic acid and the corresponding morpholines. After LC-MS confirmation, compounds were extracted with ethyl acetate or dichloromethane and purified by reverse phase HPLC.

## Acknowledgements

We thank Catherine Jones for editorial support, and Angela DeLucia, Stacey Tiamfook, JoAnn Dzink-Fox, John Walker, Whitney Barnes, Javier de Vicente, and Colin Skepper for experimental support. We also thank David Six, Charles Dean, Laura McDowell, Jennifer Leeds, Folkert Reck, Johanna Jansen, Isabel Zaror, Dirk Bussiere, Heinz Moser, Don Ganem, and past and present colleagues at Novartis Institutes for BioMedical Research in Emeryville, CA for helpful discussion and support.

## Author contributions

TU led the LpxA program. GDP and CJB performed compound selection from bacterial growth inhibition hits for target identification. RP, FC, AR, XS, and TU generated the genetic data. BCL, CMH, JS, and ML prepared the proteins and the acyl-ACP substrate. HM, DR, SM, WS, CMBR, and AB developed the biochemical assay and generated the biochemical data. MS and CW generated the SPR data. FW, AF, AOF, and AL generated the NMR data. SS, EO, and XM generated the X-ray crystallography data. WH led the chemistry efforts. PL and WH performed the inhibitor design. SS and WH synthesized the compounds. MKC and WH validated the compounds. KRP, JV, RP, and BB generated the MIC data. WH, XM, and TU wrote the draft of the paper and all authors contributed to the final manuscript.

## Data availability

Structure factors and refined atomic coordinates for the LpxA structures were deposited in the RCSB Protein Data Bank upon acceptance. The accession codes are *E. coli* LpxA in complex with compound **1**: 6P9P, and *E. coli* LpxA in complex with UDP-3-*O*-(*R*-3-hydroxymyristoyl)-GlcNAc and compound **2**: 6P9Q, compound **6**: 6P9R, compound **8**: 6P9S, and compound **13**: 6P9T. The datasets are available from the corresponding authors on reasonable request.

