## Supplemental file for "Two distinct mechanisms of small molecule inhibition of LpxA acyltransferase essential for lipopolysaccharide biosynthesis"

**This PDF file includes:**

Supplementary Methods
Figs. S1 to S5
Tables S1 to S3

**Supplementary Information Text**

**Bacterial strains, plasmids, routine growth condition, and compound susceptibility test**

Bacterial strains and plasmids used in this study are listed in Table S1. Miller's lysogeny broth (LB) [1] and LB-agar were used for routine growth and bacterial selection during plasmid construction. To maintain plasmids, 100 µg/mL ampicillin (Ap), 10 µg/mL chloramphenicol (Cm), and 50 µg/mL kanamycin (Km) were added to the media as needed. Compound susceptibility testing to determine MICs was performed in cation adjusted Mueller-Hinton II broth (CAMHB) according to CLSI protocols, as described previously [2]. To induce genes under the control of a *lac* promoter, 100 µM isopropyl β-D-1-thiogalactopyranoside (IPTG) was added to the media used in susceptibility testing. Compounds were synthesized at Novartis as described below. CHIR-090 was purchased from ChemScene (CS-0973).

**Spontaneous mutant selection**

For mutant selection with compound **1**, cells in overnight culture of MG1655 Δ*tolC* were spread on LB agar containing 0.5 µg/mL compound **1**. For mutant selection with compound **14**, overnight culture of CDY0154 was used and the cells were spread on L-agar containing 16 and 32 µg/mL compound **14**. Serial dilutions of the overnight culture were spread on LB agar to determine the number of cells (approximately $10^8$ and $10^9$ cells) used for selection. The LB agar plates were incubated overnight at 37 °C. Colonies arose and were streaked on fresh LB agar. After incubation overnight at 37 °C, a single colony of each mutant was grown in LB and stored in 15% glycerol at -80 °C for genome sequencing and susceptibility tests. The *fabZ* mutants were isolated from selection with inhibitors of LpxA and LpxD.

**Genome sequencing and mutation confirmation**

Genomic DNA for whole genome sequencing was isolated from *E. coli* strains using the Qiagen DNeasy Blood and Tissue kit with following the instruction for Gram-negative bacterial DNA. To sequence the genomes of isolated mutants, library preparation and sequencing reaction for next generation sequencing, and data analysis were performed using the Illumina Sample Preparation kits and Illumina Sequencer as described previously [2]. Sequencing reads were aligned to the reference *E. coli* MG1655 genome (Genbank ID: NC_000913.3) using bwa for determining single nucleotide polymorphisms and short insertion/deletions (indels, typically <10 bp). Mutations identified by genomics were confirmed by targeted PCR and Sanger sequencing. The primers were used to amplify the gene of interest with Phusion High-Fidelity PCR Master Mix with GC Buffer according to established protocols. Sequencing was performed by Elim BioPharma (Hayward, CA). The primers used for PCR and sequencing were TU117, TU243, TU276, TU332, TU475, TU503, TU504, TU512, and TU513.

**Construction of CDY0154**

The *acrAB* locus was replaced with the kanamycin resistant cassette ($Km^r$) using the λRED recombination system as described previously [3]. The DNA fragment to make *acrAB*::$Km^r$ was amplified from pKD13 using primers E700 and E707. The PCR product was used to transform CDY0141 carrying pKD46. The replacement of *acrAB* with $Km^r$ was confirmed with PCR using primers E466 and E702. This generated an *E. coli* strain lacking the nine efflux pumps, CDY0154 (*ΔacrAB::Km$^r$ ΔmdtBC ΔmdtF ΔmacB ΔentS ΔemrY ΔemrB ΔacrF ΔacrD*).

**Growth conditions and pooling of the *E. coli* mobile plasmid collection**

The *E. coli* mobile plasmid collection obtained from National BioResource Project in Japan is a complete set of mobile plasmid clones of *E. coli* open reading frames [4]. The expression of each open reading frame is controlled by a $P_{tac}$ promoter and LacI encoded on the same plasmid, so that gene expression is inducible by isopropyl β-D-1-thiogalactopyranoside (IPTG). Every plasmid is in an $F^+$ *recA* strain (JA200) so that it can be transferred to $F^-$ strains of *E. coli* via mating. A colony from each of the 4229 strains comprising the collection was individually inoculated in 1 mL LB supplemented with 100 µg/mL ampicillin in 96-deep-well plates. The plates were covered with breathable sealing tape and incubated at 37 °C overnight with shaking at 700 rpm. Using the overnight cultures, a pool of the entire culture collection was made. The pooled culture was stored at -80 °C in aliquots of 100 µL containing 15% glycerol. The colony forming units (CFU) of the glycerol stock was determined to be 1 x $10^8$ CFU per mL.

**Transferring of the mobile plasmid collection to NB27079-CDY0154**

One hundred microliters of the pooled culture was mixed with 100 µL overnight culture of CDY0154. Cells in the mixed culture were pelleted by centrifugation, resuspended in 300 µL LB, and spotted on LB agar. After the agar was incubated overnight, cells on agar were collected in 1 mL LB. Then 100 µL of 10-fold serial dilutions of the cell suspension were spread on LB agar supplemented with 50 µg/mL kanamycin and 50 µg/mL ampicillin. After overnight incubation at 37 °C, the cells grown on agar with the $10^{-3}$ dilution were collected in 30 mL LB. The cell suspension was mixed with 7.5 mL of 50% glycerol and the aliquots of 100 µL were stored at -80 °C. The glycerol stock was determined to be 4 x $10^8$ CFU per mL.

**Selection of CDY0154 mobile plasmid collection with compound 2**

Three milliliters LB agar were supplemented with various concentrations of compound **2**, 100 µM IPTG, 100 µg/mL ampicillin, and 50 µg/mL kanamycin in 6-well microplates. Two microliters of the pooled collection in CDY0154 (8 x $10^5$ cells) were spread on the agar. After the microplates were incubated overnight at 37 °C, colonies arising on agar were picked and grown in LB supplemented with 50 µg/mL kanamycin and with or without 100 µg/mL ampicillin. After the microplates were incubated overnight at 37 °C, colonies arising on agar were picked and grown overnight at 37 °C in 1 mL LB supplemented with 100 µg/mL ampicillin in 96-deep-well plates. The cultures were directly sent to Elim Biopharmaceuticals (Hayward, CA) for rolling circle amplification and DNA sequencing with SP6 universal primer. Obtained DNA sequence data were searched with BLAST.

**Construction of plasmids expressing LpxA-His6 and N-His-3C-Avi-LpxA**

The pET-LpxA plasmid carrying LpxA-His6 was synthesized by ATUM (Newark, CA). The vector is pJexpress_411 (very similar to pET24a) carrying the kanamycin resistance gene and a T7 promoter. Downstream of the T7 promoter, the plasmid carried the native coding sequence of *lpxA* from the *E. coli* MG1655 genome with the 18 bp sequence for C-terminal 6His tag.
The pET-avi-LpxA plasmid carrying N-His-3C-Avi-LpxA was synthesized by GENEWIZ (South Plainfield, NJ). The vector is pET24a carrying the kanamycin resistance gene and a T7 promoter. Downstream of the T7 promoter, the plasmid carried the native coding sequence of *lpxA* (without the first codon ATG) from the *E. coli* MG1655 genome with the 102 bp sequence encoding 9 histidine residues, a cleavage site of 3C protease, and the Avi tag at the N-terminus.

**Expression of *E. coli* LpxA-His6 protein**

BL21-AI was transformed with pET-LpxA and cells were spread on LB agar supplemented with 50 μg/mL Km. After incubation at 37 °C overnight, a colony arising on the agar was shaken overnight in LB supplemented with 50 μg/mL Km. The overnight culture was inoculated at 2% in fresh auto-induction medium supplemented with 50 μg/mL Km. The culture was shaken at 250 RPM at 37 °C for 2 h to reach $OD_{600}$ of 0.6, and 10 mL 20% arabinose was added to induce LpxA-His6. The culture was further incubated for 7 h (the final $OD_{600}$ of ~9) and the cells were harvested and stored at -80 °C. The recipe of the auto-induction medium used is described below.

**General methods of protein purification**

AKTA Avant Chromatography systems with Unicorn 6.0 software or higher (GE Healthcare) were used for protein purification. Protein samples were separated by SDS-PAGE using Invitrogen precast gel system in the 1 x Tris-Glycine Buffer. Novex Tris-Glycine Precast Gels with 4-20% gradient were routinely used and the Seeblue Plus2 or Sharp Stain Pre-stained Protein Standard was used as molecular weight markers. Gels were stained with Instant blue Coomassie stain for visualization and gel pictures were taken using Fluorchem imaging system. The molecular weight of the protein was checked by liquid chromatography electrospray ionization mass spectrometry (LC-MS).

**Purification of LpxA-His6 protein**

The frozen cell pellet was resuspended in 90 mL Ni [10] buffer (20 mM Tris-HCl pH 7.5, 0.3 M NaCl, 10 mM imidazole, 1 mM TCEP), and one EDTA-free protease inhibitor tablet and 10 μL of Universal nuclease (100U/mL) were added. The cells were lysed with sonication followed by centrifugation for 30 min at 50,000 x *g* at 4 °C. The supernatant was loaded onto a 5 mL Histrap FF column that was pre-equilibrated with Ni [10] buffer at 5 mL/min and washed with Ni [10] buffer until baseline. *E. coli* LpxA-His6 protein was then eluted with step gradient Ni [1000] buffer (20 mM Tris-HCl pH 7.5, 0.3 M NaCl, 1000 mM imidazole, 1 mM TCEP). Fractions were immediately analyzed by SDS-PAGE. Based on SDS-PAGE analysis, fractions containing LpxA-His6 were pooled, concentrated to 9 mL using 30k MWCO Amicon ultra (Millipore-Sigma), and loaded onto a Superdex 200 26/60 size exclusion column (GE Healthcare) that was pre-equilibrated with SEC buffer (20 mM Tris-HCl pH 7.5, 150 mM NaCl, 1 mM TCEP) at 2.5 mL/min. Elution fractions were immediately analyzed by SDS-PAGE. Fractions containing LpxA-His6 (confirmed with LC-MS) were pooled and concentrated to ~4 mL using 100 k MWCO Amicon ultra. The final protein solution (~30 mg/mL) was aliquoted and stored at -80 °C.

**Acylation of ACP and purification of acyl-ACP**

Purification of *E. coli* holo-ACP and *Vibrio harveyi* acyl-ACP synthase AaaS was performed as described previously [5]. The acylation of holo-ACP by AasS was performed by incubation for 1 h at 30 °C in a final volume of 85 mL containing of 100 mM Tris-HCl pH 7.5, 5 mM $MgCl_2$, 5 mM ATP, 0.015% Triton X-100, 1 mM TCEP, 1.17 mM DTT, 150 µM *R*-3-hydroxymyristate (50 mM stock in ethanol, Wako chemicals), 40 µM holo-ACP, and 163 nM AaaS. The reaction sample was directly loaded on to a HiTrapQ column (5 mL) pre-equilibrated with AIEX Buffer A (ABA, 20 mM Hepes-NaOH pH 8.0). The column was washed with ABA and 10% AIEX Buffer B (ABB, 20 mM Hepes-NaOH pH 8.0, 1M NaCl) in ABA. Acylated ACP was eluted with a linear gradient of 10 to 70% ABB over 100 mL. Fractions were analyzed with SDS-AGE and LC-MS. Fractions containing acyl-ACP were pooled, concentrated using an Amicon Ultra-4 centrifugal filter unit MWCO 3000 to 30 mg protein/mL, and loaded on to a HiLoad 16/60 Superdex 75 prep grade column pre-equilibrated with 20 mM Hepes-NaOH pH 8.0 containing 150 mM NaCl and 1 mM TCEP. The protein was eluted in the same buffer at a flow rate of 1 mL/min. Fractions were analyzed with SDS-AGE and LC-MS. Fractions containing acyl-ACP were polled and concentrated to approximately 30 mg/mL as described above.

**Expression of N-His-3C-Avi-LpxA and preparation of biotinylated Avi-LpxA**

BL21(DE3) was transformed with pET-avi-LpxA and cells were spread on LB agar supplemented with 50 µg/mL Km. After incubation at overnight, colonies arising on the agar was suspended in 100 mL Terrific Broth supplemented with 50 mM MOPS-NaOH pH 7.5 and 50 µg/mL Km. The culture was shaken at 250 RPM overnight. The next day, 30 mL of the overnight culture was inoculated into 3L TB supplemented with 50 mM MOPS-NaOH pH 7.5 and 50 µg/mL Kan. After the culture was shaken at 250 RPM to reach $OD_{600}$ of 0.7, 1 mM IPTG was added to induce N-His-3C-Avi-LpxA. The culture was further incubated at 25 °C overnight. The cells were harvested with centrifugation (5000 x *g*, 30 min, 4 °C) and 10-15 g of cell biomass was stored in each 50 mL conical tube at -80 °C.

Frozen cell biomass (10-15 g) overexpressing N-His-3C-Avi-LpxA was resuspended in 40 mL Ni-A buffer (50 mM HEPES-NaOH pH 8.0, 0.5 M NaCl, 10% glycerol, 1 mM TCEP) supplemented with 50 µL Promega Protease Inhibitor Cocktail and 10,000U Universal nuclease. The cells were lysed with sonication on ice for 2 min (10 sec burst, 30 sec rest, 70% output) followed by centrifugation at 40,000 x *g* for 30 min at 4 °C. The supernatant was loaded onto a 5 mL His-Trap FF column that was pre-equilibrated with Ni-A buffer, and washed with Ni-A buffer until baseline. N-His-3C-Avi-LpxA protein was then eluted with step gradient of 10% and 80% Ni-B buffer (Ni-A buffer with 500 mM imidazole). Fractions were analyzed by SDS-PAGE and fractions containing N-His-3C-Avi-EcLpxA were pooled (92.6 mg protein in 63 mL). To generate Avi-LpxA, the pooled eluent was incubated overnight at 4 °C with 30,000 U of 3C protease during dialysis against 50 mM HEPES-NaOH buffer at pH 8.0 containing 150 mM NaCl and 1 mM TCEP. The majority of the protein was lost due to precipitation under the low-salt condition, which was required for efficient biotinylation. To stabilize the remaining protein, additional NaCl was added to the sample for a final NaCl concentration of 250 mM. The precipitated protein was removed by centrifugation at 4,000 x *g* for 10 min. The Avi-EcLpxA protein was biotinylated in vitro by BirA biotin-protein ligase. The biotinylation reaction was performed with incubation overnight at 4 °C in a 70 mL sample containing Avi-EcLpxA, 200 ng BirA 10 mM ATP, 10 mM $MgCl_2$, 150 µM biotin. LC-MS analysis confirmed 100% 3C cleavage and >95% biotinylation.

The sample was then loaded on a His-Trap FF column and proteins were eluted with step gradient as described above. Fractions were analyzed by SDS-PAGE. Flow-through fractions containing biotinylated Avi-EcLpxA were pooled, and concentrated using an Amicon 50 kDa MWCO concentrator to 5.5 mL, and loaded onto a Superdex 200 16/60 size exclusion column that was pre-equilibrated with SEC buffer (20 mM HEPES-NaOH pH 8.0, 300 mM NaCl, 1 mM TCEP). Elution fractions were collected at 1 mL/min and analyzed by SDS-PAGE. The chromatogram traces showed a high A260/A280 ratio, indicating the presence of excess ATP in the protein fractions. The fractions containing the protein were pooled, diluted, and loaded on to a HiTrap Q column which was pre-equilibrated with ANX-A buffer (50 mM HEPES-NaOH pH 8.0, 50 mM NaCl, 1 mM TCEP). The protein was eluted with a gradient 0-70% of ANX-B (50 mM HEPES-NaOH pH 8.0, 1000 mM NaCl, 1 mM TCEP). Fractions containing the protein were pooled, concentrated using an Amicon 50 kDa MWCO concentrator to 5 mL, and loaded again onto a Superdex 200 16/60 size exclusion column that was pre-equilibrated with SEC buffer (20 mM HEPES pH 8.0, 300 mM NaCl, 1 mM TCEP). Fractions containing biotinyalated Avi-LpxA (~40 mL) were pooled and concentrated to 0.75 mL using an Amicon 50kDa MWCO concentrator. The final protein solution (3.4 mg/mL) was aliquoted and stored at -80 °C.

**Preparation of isotope-enriched LpxA-His6 protein for NMR studies**

For NMR experiments, uniformly $^{15}N/^{2}H$ labeled LxpA-His6 protein with selectively introduced $^{13}C$ methyl groups was prepared. To maximize the number of NMR-active probes, we simultaneously labeled the methyl groups of all methyl containing amino acids (methionine, isoleucine, leucine, valine, alanine, threonine, known as MILVAT labeling scheme) according to the recently published protocol [6-7]. Purification of isotope labeled LpxA-His6 protein was performed as described above for unlabeled LpxA-His6 protein, except that buffer comprised solely of 50 mM sodium phosphate monobasic pH 7.0 was used. The purified protein solution was concentrated and stored at -80 °C.

#### LpxA SPE-MS biochemical/enzyme assay and $IC_{50}$ determination

The LpxA enzymatic reaction was performed in a 384-well plate in a final volume of 10 µL 50 mM sodium phosphate buffer (pH 7) containing 100 mM NaCl, 0.01% Tween-20, 1 mM TCEP, 1 mM EDTA, 0.01% BSA, 1 nM LpxA-His6, 1 µM *R*-3-hydroxymyristate-ACP, 200 µM UDP-GlcNAc, and test compound (8 points, 3-fold dilutions, top concentration: 200 µM). The stock solution of each compound was made in 10 mM in DMSO. The concentration range was between 0.114 µM to 250 µM or between 1.7 nM to 250 µM (8-point or 16-point, 3× dilution series). The concentrations of *R*-3-hydroxymyristate-ACP and UDP-GlcNAc were the apparent $K_M$ values of the substrates. After the reaction was incubated for 15 min at room temperature, the reaction was quenched by addition of 90 µL 2.22% acetic acid (final concentration, 2% acetic acid). The generated LpxA product UDP-3-*O*-*R*-3-hydroxymyristate-GlcNAc (*m/z* = 832.3) was detected with a SPE-MS system using Agilent RapidFire 365 coupled to a Sciex 5500 triple quadrupole MS with a Turbo V ESI source with product ions at: 158.9, 273.0, and 385.0 *m/z*. The SPE was performed using a reversed-phase Agilent Type A (C4) cartridge with a setting of pump 1 at a flow-rate of 1.5 mL/min of mobile phase A (10 mM ammonium acetate in water) and pump 2 at a flow-rate of 1.25 mL/min of mobile phase B (25% acetone, 25% acetonitrile, and 50% water). The MS settings were: Curtain gas = 20 psig, Collision gas = 12 psig, IonSpray Voltage = -4500V, temperature = 650°C, Ion source gas 1 = 50 psig, Ion source gas 2 = 50 psig. Data were analyzed with Agilent RapidFire Integrator software. Area under the curve results were then used to calculate percent inhibition and $IC_{50}$ values were determined as described previously [8]. Mean ± standard error of the mean (SEM) or standard deviation of the mean (SD) values were calculated using Microsoft Excel 2010 software. In this assay, the $IC_{50}$ value of peptide 920, the known peptide inhibitor of LpxA [9-10], as determined to be 1.9 µM ± 0.8 µM, which reasonably matched the published value of 4.7 ± 0.2 µM obtained using the ThioGlo detection method [11].

#### Surface plasmon resonance binding assay

The SPR method to analyze small molecule interactions was developed and reported previously [12]. The instrument used for these studies was Biacore T200. Software used to perform the SPR assay was Biacore T200 Control Software v2.0. The instrument was primed with SPR buffer (50 mM HEPES pH 8.0, 150 mM NaCl, 1 mM TCEP, 2 mM EDTA, 0.05% P20). It should be noted that biotinylated Avi-LpxA was pH sensitive, and this assay did not work well at pH 7. A streptavidin-coated SPR chip was inserted into the instrument and primed. The biotinylated *E. coli* LpxA protein was immobilized onto a chip flow cell. Start-up cycles (10 cycles) were run followed by an 11-point DMSO standard curve (typically 1-3% DMSO). The N-terminal biotinylated *E. coli* LpxA protein was used because it generated higher signals of ligand binding than C-terminal biotinylated LpxA. Sample analysis occurred at a flow rate of 100 µL/min at 20°C with a 30 second injection and a 60 second dissociation. Dilutions of compound were prepared by mixing 2 µL DMSO stock solution with 98 µL SPR buffer in a deep-well 384-well plate. Data were analyzed using Evaluation Software v2.0.

**Protein observed 2D NMR assay**

All protein-observed NMR spectra were acquired at 305 K on a Bruker AVANCE-500 spectrometer, equipped with a 1.7 mm TCI probe and controlled by Topspin 3.5 pl7 software (Bruker). Samples contained 80 µM isotope-labeled (MILVAT methyl labeled) LpxA, and 100 µM compound **1** or compound **2** with 150 µM UDP-3-*O*-(*R*-3-hydroxymyristoyl)-GlcNAc (Toronto Research Chemicals) in 50 mM sodium phosphate buffer pH 7.0, 100 mM sodium chloride, 2 mM deuterated DTT, and 10 µM DSS. Two-dimensional $^{13}C$-$^{1}H$ SOFAST HMQC spectra [6-7, 13] were recorded with 64 scans and up to 96 points in the indirect dimension.

**Crystallization and structure determination**

The concentration of the LpxA-His6 protein was adjusted to 15 mg/mL in 20 mM Tris-HCl pH 7.5, 150 mM NaCl, 1 mM TCEP. LpxA-His6 (~15 mg/mL) was crystallized in 0.1 M tri-sodium citrate pH 4.4 - 4.7, 7-9% PEG1000, and using 1:1 ratio of protein to crystallant by the sitting-drop vapor diffusion method. Compound **1** was supplemented to the soaking buffer (0.1M Na/K phosphate pH 7, 10% PEG1000) at 2 mM final concentration. Compound **2** and UDP-3-*O*-(*R*-3-hydroxymyristoyl)-GlcNAc were supplemented at 2 mM and 40 mM final concentration, respectively. Before data collection, crystals were cryoprotected in soaking buffer plus 20% glycerol and flash cooled in liquid nitrogen. Data were collected on a single crystal cooled to 100K using an R-AXIS detector (Rigaku) and in-house Cu Kα X-ray source (FR-E SuperBright High-Brilliance Rotating Anode Generator). Data were processed using autoPROC (Global Phasing, LTD). The structures of *E. coli* LpxA in complex with ligands were solved by molecular replacement with PHASER [14] using the published crystal structure of the binary complex of *E.coli* LpxA/LpxA product (PDB ID code 2QIA) [15] as the starting model. Compound structures were first placed by the rhofit (Global Phasing, LTD) and further refined in Phenix [16]. Subsequent rounds of model building and refinement with Phenix.refine program were carried out until convergence.

**Chemical synthesis**

Compound **1** and 5-(4-methylpyridin-2-yl)-1H-pyrazole-3-carboxylic acid for compound **2** were prepared by literature procedures [17-18]. When morpholine intermediates were not commercially available, they were prepared by two different methods [19-20]. Compound **2** and the analogs were prepared by amide coupling between 5-(4-methylpyridin-2-yl)-1H-pyrazole-3-carboxylic acid and the corresponding morpholines as described below. After LC-MS confirmation, compounds were extracted with ethyl acetate (EtOAc) or dichloromethane (DCM) and purified by reverse phase

HPLC. When a racemic mixture was synthesized (compounds **9** and **10**), two enantiomers were resolved by chiral supercritical fluid chromatography (SFC). The absolute configurations were assigned on the basis of biochemical $IC_{50}$ values and X-ray structural information. The resolution of a racemic mixture of compound **13** and its enantiomer under several chiral separation conditions failed. The absolute configuration of compound **13** was assigned on the basis of $^{1}$H-$^{13}$C HMQC results of (*R*)- or (*S*)-(3-bromo-1H-pyrazol-5-yl)(3-(2-chloro-6-methoxybenzyl)morpholino)methanone in the presence of LpxA product complex. The chemical structures of synthesized compounds were confirmed by $^{1}$H and/or $^{13}$C NMR spectroscopy and high resolution LC-MS. The rotational barrier of the amide bond of compound **6** was determined by variable-temperature NMR.

**Synthesis of 3-(2-(4-methoxyphenyl)-2-oxoethyl)-5,5-diphenylimidazolidine-2,4-dione (compound 1)**

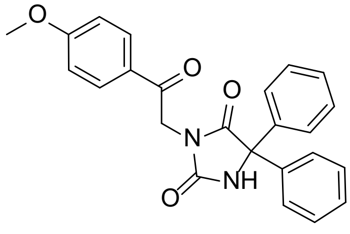

compound **1**

LC-MS (m/z): 401.1 $[M+H]^+$. HRMS (ESI) *m/z* calculated for C24H20N2O4 400.1423 $[M+H]^+$ found 401.1487. $^{1}$H NMR (600 MHz, DMSO-*d*$^{6}$) δ 9.72 (s, 1H), 8.04 (d, *J* = 9.0 Hz, 2H), 7.46 – 7.41 (m, 8H), 7.41 – 7.36 (m, 2H), 7.10 (d, *J* = 8.9 Hz, 2H), 4.99 (s, 2H), 3.87 (s, 3H). $^{13}$C NMR (151 MHz, DMSO-*d*$^{6}$) δ 190.25, 173.42, 163.85, 154.94, 139.55, 130.54, 128.48, 128.19, 126.88, 119.48, 114.19, 69.63, 55.63, 44.48.

**Synthesis of morpholine intermediates**

**Method 1**[19]

**(*R*)-3-(2-fluorobenzyl)morpholine**

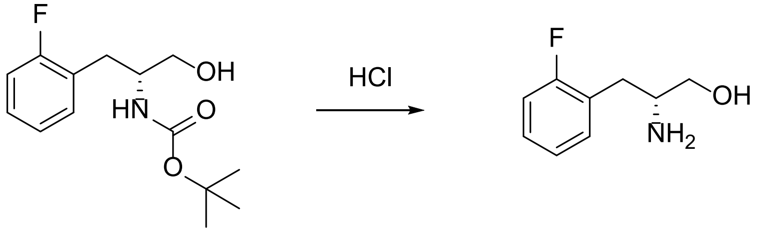

<u>tert-butyl (*R*)-(1-(2-fluorophenyl)-3-hydroxypropan-2-yl)carbamate</u>

To a solution of (*R*)-2-amino-3-(2-fluorophenyl)propan-1-ol (2.95 g, 10.97 mmol) in DCM (10 mL) was added 4 M HCl in dioxane (8 mL, 32 mmol) at 0 °C. The reaction mixture was stirred at room temperature for 1 h. LC-MS indicated that no starting materials remained. The volatile materials were removed in vacuo. The crude HCl salt of (*R*)-2-amino-3-(2-fluorophenyl)propan-1-ol was used in the next step without further purification. LC-MS (m/z): 170.1 $[M+H]^+$.

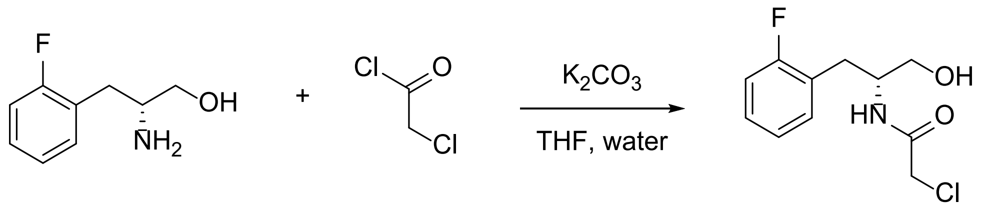

(*R*)-2-chloro-N-(1-(2-fluorophenyl)-3-hydroxypropan-2-yl)acetamide

To a solution of (*R*)-2-amino-3-(2-fluorophenyl)propan-1-ol (2 g, 9.72 mmol) in THF (20 mL) and water (20 mL) was added potassium carbonate (4.03 g, 29.2 mmol) followed by slow addition of 2-chloroacetyl chloride (0.78 mL, 9.72 mmol) at 0 °C. The reaction mixture was stirred at room temperature for 2 h. LC-MS indicated that no starting materials remained. The reaction mixture was extracted with EtOAc (50 mL × 2). The combined organic layers were dried over anhydrous sodium sulfate, filtered off, and concentrated in vacuo. The crude product was purified by ISCO (0 to 100% EtOAc in heptane) to yield (*R*)-2-chloro-N-(1-(2-fluorophenyl)-3-hydroxypropan-2-yl)acetamide (1.12 g, 4.56 mmol, 47% yield). LC-MS (m/z): 246.0 $[M+H]^+$.

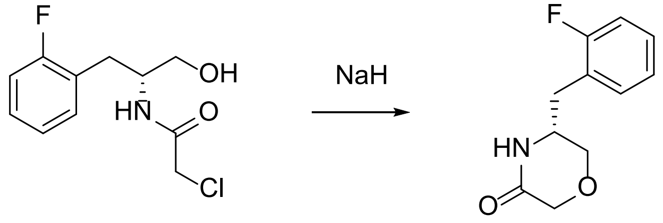

(*R*)-5-(2-fluorobenzyl)morpholin-3-one

To a solution of (*R*)-2-chloro-N-(1-(2-fluorophenyl)-3-hydroxypropan-2-yl)acetamide (1.463 g, 5.95 mmol) in THF (20 mL) was added sodium hydride (60% in oil, 0.476 mg, 11.91 mmol) at 0 °C. The reaction mixture was stirred at room temperature for 0.5 h. LC-MS indicated that no starting materials remained. After quenched with saturated ammonium chloride solution (20 mL), the reaction mixture was extracted with EtOAc (50 mL × 2). The combined organic layers were dried over anhydrous sodium sulfate, filtered off, and concentrated in vacuo. The crude product was purified by ISCO (0 to 100% EtOAc in heptane) to yield (*R*)-5-(2-fluorobenzyl)morpholin-3-one (1.01 g, 4.83 mmol, 81% yield). LC-MS (m/z): 209.9 $[M+H]^+$.

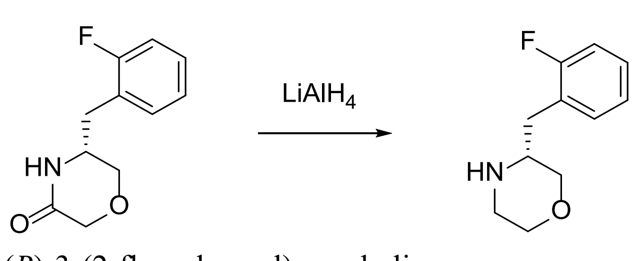

(*R*)-3-(2-fluorobenzyl)morpholine

To a solution of (*R*)-5-(2-fluorobenzyl)morpholin-3-one (1.01 g, 4.83 mmol) in THF (20 mL) was added lithium aluminum hydride (2 M solution in THP, 9.66 mL, 19.31 mmol) at 0 °C. The reaction mixture was stirred at room temperature for 16 h. LC-MS indicated that no starting materials remained. After quenched with Rochelle salt solution (20 mL), the reaction mixture was stirred for 1 h. The mixture was filtered through Celite® pad. The filtrate was extracted with EtOAc (50 mL × 2). The combined organic layers were dried over anhydrous sodium sulfate, filtered off, and concentrated in vacuo. The crude (*R*)-3-(2-fluorobenzyl)morpholine was used in the next step without further purification (0.9 g, 4.61 mmol, 95% yield). LC-MS (m/z): 196.1 $[M+H]^+$.

**(*R*)-3-(2-methoxybenzyl)morpholine**

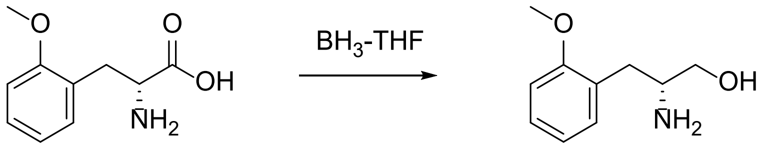

(*R*)-2-amino-3-(2-methoxyphenyl)propan-1-ol

To a solution of (*R*)-2-amino-3-(2-methoxyphenyl)propanoic acid (1 g, 5.12 mmol) was added borane tetrahydrofuran complex (15.37 ml, 15.37 mmol) at 0 °C. The mixture was stirred at room temperature for 16 h. LC-MS indicated that no starting materials remained. After the reaction mixture was quenched with MeOH (3 mL) and was stirred for 1 h, the volatile materials were removed in vacuo. To this, 3 N HCl (1 mL) and MeOH (ca. 2 mL) was added. After MeOH was evaporated, the mixture was diluted with water and extracted with DCM. The separated aqueous layers were basified by 3 M NaOH solution up to pH 10 and extracted by DCM three times. The combined organic layers were concentrated to yield (*R*)-2-amino-3-(2-methoxyphenyl)propan-1-ol (744 mg, 4.11 mmol, 80%), which was used in the next step without further purification. LC-MS (m/z): 182.2 $[M+H]^+$.

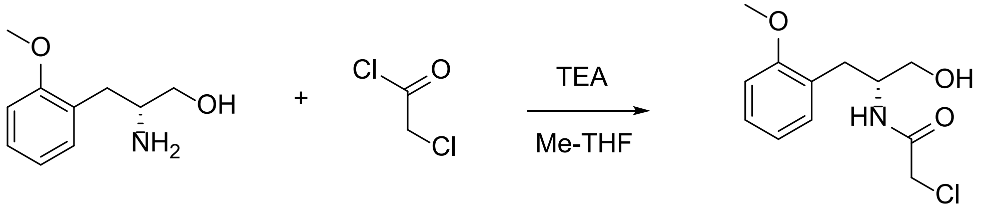

(*R*)-2-chloro-N-(1-hydroxy-3-(2-methoxyphenyl)propan-2-yl)acetamide

To a solution of (*R*)-2-amino-3-(2-methoxyphenyl)propan-1-ol (0.74 g, 4.08 mmol) in methyl-THF (13.6 mL) was slowly added 2-chloroacetyl chloride 2-chloroacetyl chloride (0.358 ml, 4.49 mmol) at 0 °C. The reaction mixture was stirred at room temperature for 1.5 h. LC-MS indicated that no starting materials remained. After adding water (20 mL), the reaction mixture was extracted with EtOAc (20 mL × 2). The combined organic layers were dried over anhydrous sodium sulfate, filtered off, and concentrated in vacuo. The crude (*R*)-2-chloro-N-(1-hydroxy-3-(2-methoxyphenyl)propan-2-yl)acetamide (0.873 g, 3.39 mmol, 83% yield) was used in the next step without further purification. LC-MS (m/z): 258.2 $[M+H]^+$.

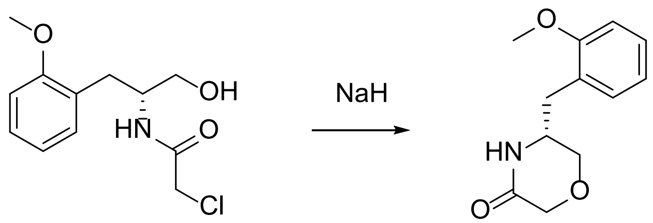

<u>(*R*)-5-(2-methoxybenzyl)morpholin-3-one</u>

To a solution of (*R*)-2-chloro-N-(1-hydroxy-3-(2-methoxyphenyl)propan-2-yl)acetamide (0.873 g, 3.39 mmol) in THF (80 mL) was added sodium hydride (60% in oil, 0.142 mg, 3.56 mmol) at 0 °C. The reaction mixture was stirred at room temperature for 0.5 h. LC-MS indicated that no starting materials remained. After quenched with saturated ammonium chloride solution (50 mL), the reaction mixture was extracted with EtOAc (50 mL × 2). The combined organic layers were dried over anhydrous sodium sulfate, filtered off, and concentrated in vacuo. The crude (*R*)-5-(2-methoxybenzyl)morpholin-3-one was used in the next step without further purification. LC-MS (m/z): 222.2 $[M+H]^+$.

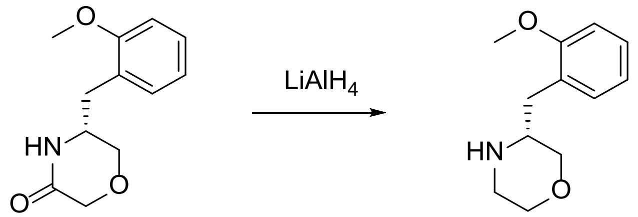

<u>(*R*)-3-(2-methoxybenzyl)morpholine</u>

To a solution of (*R*)-5-(2-methoxybenzyl)morpholin-3-one (0.749 g, 15.23 mmol) in THF (34 mL) was added lithium aluminum hydride (2 M solution in THP, 6.62 mL, 19.31 mmol) at 0 °C. The reaction mixture was stirred at room temperature for 16 h. LC-MS indicated that no starting materials remained. After quenched with Rochelle salt solution (20 mL), the reaction mixture was stirred for 1 h. The mixture was filtered through Celite® pad. The filtrate was extracted with EtOAc (50 mL × 2). The combined organic layers were dried over anhydrous sodium sulfate, filtered off, and concentrated in vacuo. The crude (*R*)-3-(2-methoxybenzyl)morpholine was used in the next step without further purification (0.7 g, 3.39 mmol, >99% yield). LC-MS (m/z): 208.3 $[M+H]^+$.

**(*R*)-3-(2-chlorobenzyl)morpholine**

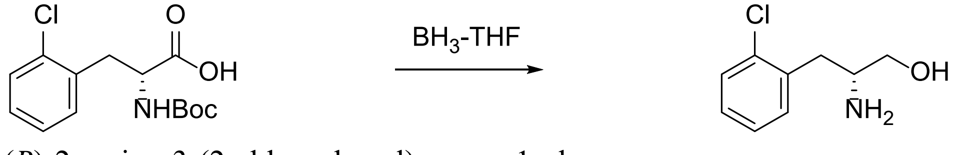

<u>(*R*)-2-amino-3-(2-chlorophenyl)propan-1-ol</u>

To a solution of (*R*)-2-((tert-butoxycarbonyl)amino)-3-(2-chlorophenyl)propanoic acid (1 g, 3.34 mmol) in THF (11 mL) was added borane tetrahydrofuran complex (1 M THF solution, 10.01 mL, 10.01 mmol) at 0 °C. After the mixture was stirred at room temperature for 16 h, the reaction mixture was quenched with MeOH (5 mL) and the reaction mixture was stirred for 1 h. The volatile materials were removed in vacuo. To this, 3 N HCl aqueous solution (1 mL) and MeOH (ca. 3 mL) was added. The mixture was stirred for 1 h. After MeOH was evaporated, the mixture was extracted with diethyl ether (10 mL). The separated aqueous layer was basified by 3 M NaOH solution up to pH 10 and extracted by DCM three times. The organic layers were concentrated to provide (*R*)-2-amino-3-(2-chlorophenyl)propan-1-ol, which was used in the next step without further purification. LC-MS (m/z): 186 [M+H-Boc]$^+$.

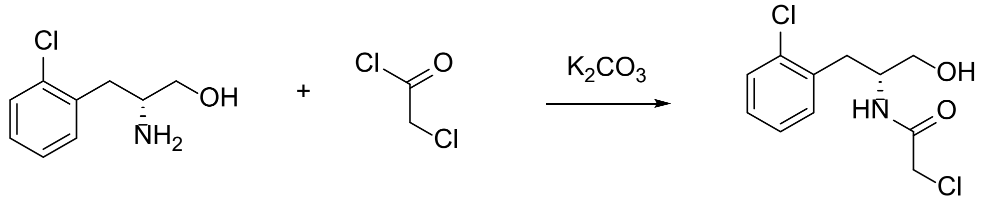

(*R*)-2-chloro-*N*-(1-(2-chlorophenyl)-3-hydroxypropan-2-yl)acetamide

To a solution of (*R*)-2-amino-3-(2-chlorophenyl)propan-1-ol (0.61 g, 2.04 mmol) in THF (10 mL) and water (10 mL) was added potassium carbonate (0.844 g, 6.11 mmol) followed by slow addition of 2-chloroacetyl chloride (0.18 mL, 2.24 mmol) at 0 °C. The reaction mixture was stirred at room temperature for 2 h. LC-MS indicated that no starting materials remained. The reaction mixture was extracted with EtOAc (20 mL × 2). The combined organic layers were dried over anhydrous sodium sulfate, filtered off, and concentrated in vacuo to yield crude (*R*)-2-chloro-N-(1-(2-chlorophenyl)-3-hydroxypropan-2-yl)acetamide (0.53 g, 2.02 mmol, 99% yield). The crude product was used in the next step without further purification. LC-MS (m/z): 262.0 [M+H]$^+$.

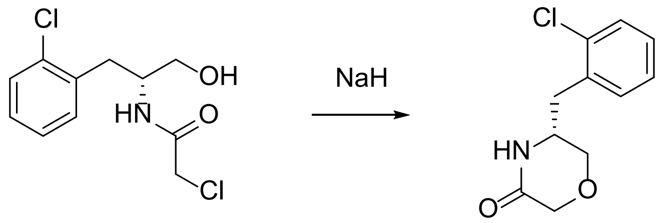

(*R*)-5-(2-chlorobenzyl)morpholin-3-one

To a solution of (*R*)-2-chloro-N-(1-(2-chlorophenyl)-3-hydroxypropan-2-yl)acetamide (0.53 g, 2.02 mmol) in THF (6.7 mL) was added sodium hydride (60% in oil, 0.24 mg, 6.07 mmol) at 0 °C. The reaction mixture was stirred at room temperature for 0.5 h. LC-MS indicated that no starting materials remained. After quenched with saturated ammonium chloride solution (10 mL), the reaction mixture was extracted with EtOAc (30 mL × 2). The combined organic layers were dried over anhydrous sodium sulfate, filtered off, and concentrated in vacuo to yield crude (*R*)-5-(2-chlorobenzyl)morpholin-3-one (0.45 g, 2.0 mmol, 99% yield). LC-MS (m/z): 249 [M+H]$^+$.

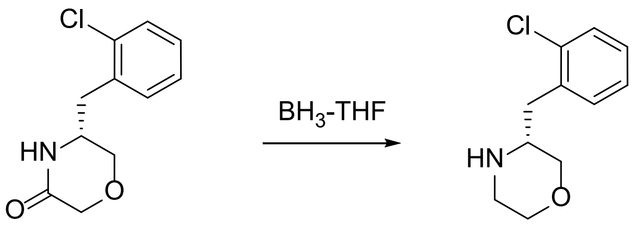

(*R*)-3-(2-chlorobenzyl)morpholine

To a solution of (*R*)-5-(2-chlorobenzyl)morpholin-3-one (0.45 g, 1.994 mmol) in THF (6.6 mL) was added borane tetrahydrofuran complex (1 M THF solution, 5.98 mL, 5.98 mmol) at 0 °C. After the mixture was stirred at room temperature for 16 h, the reaction mixture was quenched with MeOH (5 mL) and was stirred for 1 h. After the volatile materials were removed in vacuo, the mixture was diluted with EtOAc (5 mL) and water (5 mL). The mixture was extracted with EtOAc. The combined organic layers were dried over anhydrous sodium sulfate, filtered off, and concentrated in vacuo to yield crude (*R*)-3-(2-chlorobenzyl)morpholine (0.32 g, 1.5 mmol, 76%). The crude product was used in the next step without further purification. LC-MS (m/z): 212 $[M+H]^+$.

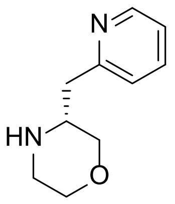

**(*R*)-3-(pyridin-2-ylmethyl)morpholine** was prepared by method 1 using commercially available (*R*)-2-amino-3-(pyridin-2-yl)propanoic acid. LC-MS (m/z): 179.9 $[M+H]^+$.

**Method 2**[20]

**(+/-)-3-(2-fluoro-6-methoxybenzyl)morpholine**

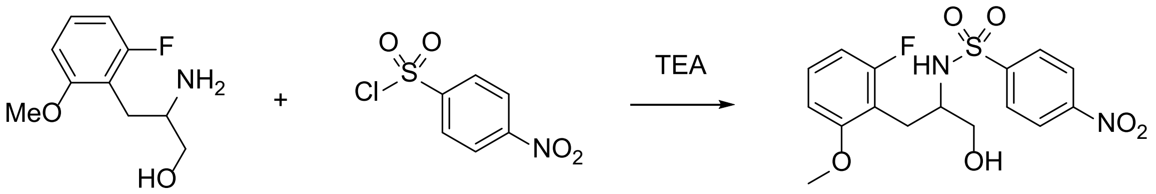

(+/-)-N-(1-(2-fluoro-6-methoxyphenyl)-3-hydroxypropan-2-yl)-4-nitrobenzenesulfonamide

To a solution of 2-amino-3-(2-fluoro-6-methoxyphenyl)propan-1-ol (315 mg, 1.581 mmol) in DCM (5 mL) was added TEA (331 µl, 2.372 mmol) and 4-nitrobenzene-1-sulfonyl chloride (368 mg, 1.66 mmol). The reaction mixture was stirred at room temperature for 3 h. After quenched with water (5 mL), the organic layers were separated and the aqueous layers were extracted with DCM (5 mL × 2). The combined organic layers were washed with water and brine, dried over anhydrous sodium sulfate, filtered off, and concentrated in vacuo. The crude product was purified by ISCO (0 to 30% EtOAc in heptane) to yield (+/-)-N-(1-(2-fluoro-6-methoxyphenyl)-3-hydroxypropan-2-yl)-4-nitrobenzenesulfonamide (215 mg, 0.56 mmol, 35% yield). LC-MS (m/z): 385.0 $[M+H]^+$.

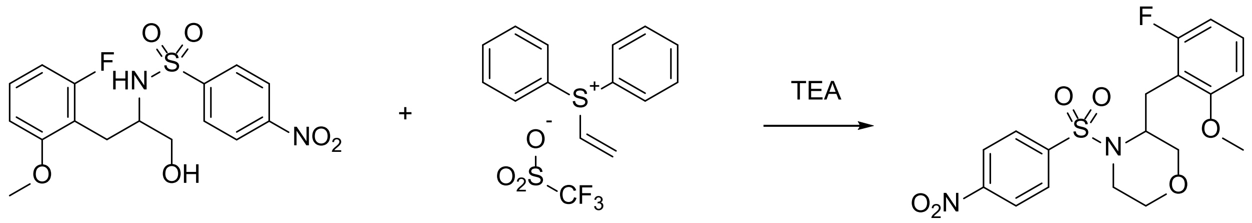

(+/-)-3-(2-fluoro-6-methoxybenzyl)-4-((4-nitrophenyl)sulfonyl)morpholine

To a solution of N-(1-(2-fluoro-6-methoxyphenyl)-3-hydroxypropan-2-yl)-4-nitrobenzenesulfonamide (215 mg, 0.559 mmol) in DCM (1.86 mL) in a microwave vessel was added triethylamine (312 µl, 2.237 mmol) followed by slow addition of diphenyl(vinyl)sulfonium trifluoromethanesulfonate (304 mg, 0.839 mmol) in DCM (1.86 mL) at 0 °C. After the reaction mixture was stirred at 0 °C for 1 h, the reaction mixture was heated to 90 °C for 45 min at a microwave reactor. After quenched with 1 M sodium phosphate monobasic solution (3 mL), the organic layers were separated and washed with water and brine, dried over anhydrous sodium sulfate, filtered off, and concentrated in vacuo. The crude product was purified by ISCO (0 to 30% EtOAc in heptane) to yield (+/-)-3-(2-fluoro-6-methoxybenzyl)-4-((4-nitrophenyl)sulfonyl)morpholine (158 mg, 0.385 mmol, 68.8 % yield). LC-MS (m/z): 597.2 $[M+H]^+$.

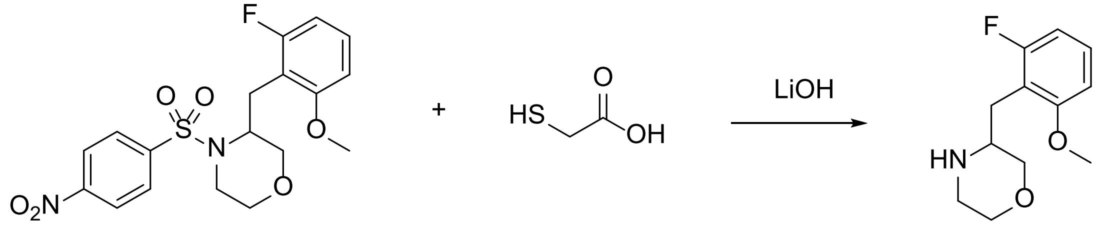

(+/-)-3-(2-fluoro-6-methoxybenzyl)morpholine

To a stirred suspension of lithium hydroxide (97 mg, 2.310 mmol) monohydrate in DMSO (1.3 mL) under nitrogen was added 2-mercaptoaceticacid (53.5 µl, 0.770 mmol). The reaction mixture was stirred for 5 min at room temperature. To this mixture was added 3-(2-fluoro-6-methoxybenzyl)-4-((4-nitrophenyl)sulfonyl)morpholine (158 mg, 0.385 mmol) in DMSO (1 mL). The reaction mixture was stirred for 16 h. After quenched with 2 M potassium carbonate solution, the organic layers were separated and the aqueous layers were extracted with DCM (2 mL × 2). The combined organic layers were washed with water twice and then dried over anhydrous sodium sulfate, filtered off, and concentrated in vacuo. The crude (+/-)-3-(2-fluoro-6-methoxybenzyl)morpholine was used in the next step without further purification (86 mg, 0.37 mmol, >99 % yield). LC-MS (m/z): 226.2 $[M+H]^+$.

**Method 3**

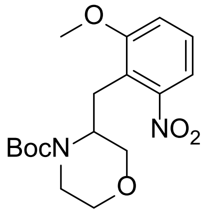

**(+/-)-tert-butyl 3-(2-methoxy-6-nitrobenzyl)morpholine-4-carboxylate** was synthesized from a custom synthesis provider. LC-MS (m/z): 374.9 $[M+Na]^+$. $^1$H NMR (300 MHz, DMSO-$d^6$) δ 8.47- 7.34 (m, 3 H), 4.21- 4.20 (m, 1 H), 3.85- 3.73 (m, 5 H), 3.64-3.49 (m, 2 H), 3.34- 3.32 (m, 2 H), 3.16- 3.12 (m, 1 H), 2.72- 2.70 (m, 1 H), 1.22- 0.97 (m, 9 H).

**(+/-)-3-(2-chloro-6-methoxybenzyl)morpholine**

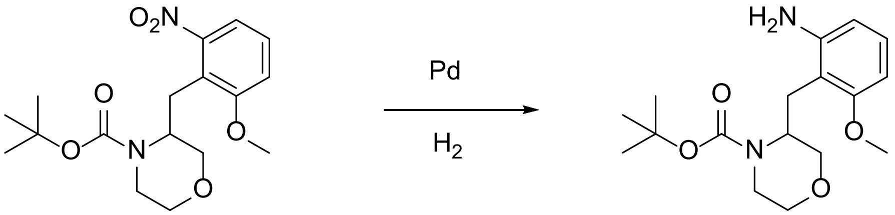

(+/-)-tert-butyl 3-(2-amino-6-methoxybenzyl)morpholine-4-carboxylate

A solution of (+/-)-tert-butyl 3-(2-methoxy-6-nitrobenzyl)morpholine-4-carboxylate (10 g, 28.4 mmol) in MeOH (70.9 mL) and EtOAc (70.9 mL) was flushed with nitrogen stream for 5 min, followed by addition of Pd-C (3.02 g, 2.84 mmol). The reaction mixture was flushed with hydrogen gas for 5 min and equipped with hydrogen gas balloon. The mixture was stirred for 16 h at room temperature. The reaction mixture was filtered off Celite® pad. The volatile filtrates were removed in vacuo to yield (+/-)-tert-butyl 3-(2-amino-6-methoxybenzyl)morpholine-4-carboxylate (9 g, 27.9 mmol, 98 % yield). LC-MS (m/z): 323.4 $[M+H]^+$.

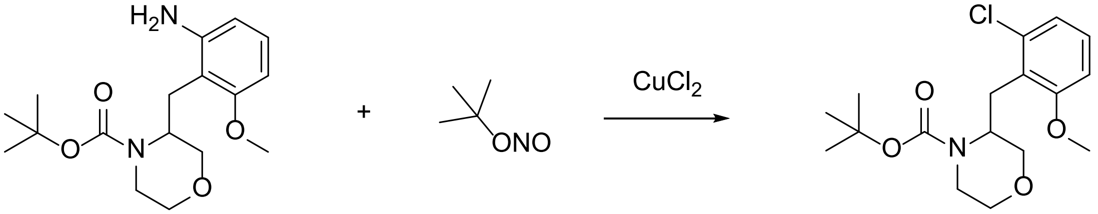

(+/-)-tert-butyl 3-(2-chloro-6-methoxybenzyl)morpholine-4-carboxylate

To a solution of tert-butyl 3-(2-amino-6-methoxybenzyl)morpholine-4-carboxylate (7 g, 21.71 mmol) and copper(II) chloride (3.50 g, 26.1 mmol) in MeCN (109 mL) was added tert-butyl nitrite (3.87 mL, 32.6 mmol). The reaction mixture was stirred at room temperature for overnight. After the mixture was quenched with sodium bicarbonate solution, the solids formed were filtered off Celite® pad. After pyridine (2 mL) was added to the filtrate, the organic layer was extracted with EtOAc. Then, the organic layers were washed with water and brine, dried over $Na_2SO_4$, filtered off and concentrated in vacuo. The crude product was purified by ISCO twice (gradient EtOAc in heptane; a dichlorinated byproduct was removed in the second purification step) to yield (+/-)-tert-butyl 3-(2-chloro-6-methoxybenzyl)morpholine-4-carboxylate (3.07 g, 8.98 mmol, 41.4% yield). LC-MS (m/z): 286.3 $[M+H]^+$. $^1$H NMR (400 MHz, $CDCl_3$) δ 7.11 (t, $J$ = 8.2 Hz, 1H), 6.97 (t, $J$ =

8.7 Hz, 1H), 6.73 (d, $J$ = 8.3 Hz, 1H), 4.34 (s, 1H), 3.91 (d, $J$ = 8.5 Hz, 1H), 3.84 (m, 1H), 3.81 (s, 3H), 3.70 – 3.38 (m, 5H), 2.79 (d, $J$ = 13.6 Hz, 1H), 1.55 (s, 9H).

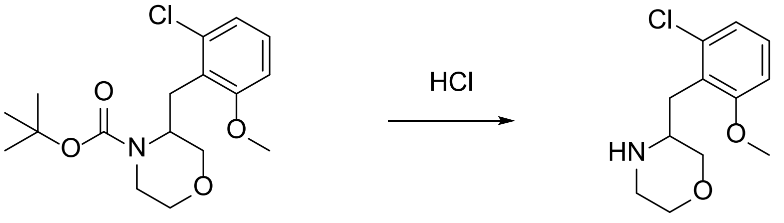

(+/-)-3-(2-chloro-6-methoxybenzyl)morpholine

To a solution of (+/-)-tert-butyl 3-(2-chloro-6-methoxybenzyl)morpholine-4-carboxylate (3.07 g, 8.98 mmol) in DCM (1 mL) was added 1 M HCl solution in dioxane (22.45 mL, 90 mmol). The reaction mixture was stirred overnight. The volatile materials were removed in vacuo. The crude product was used in the next step without further purification (2.82 g, 8.96 mmol, >99% yield). LC-MS (m/z): 244.3 $[M+3H]^+$.

**Synthesis of 5-(4-methylpyridin-2-yl)-1H-pyrazole-3-carboxylic acid[18]**

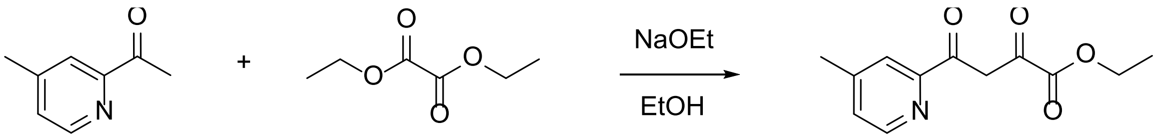

ethyl 4-(4-methylpyridin-2-yl)-2,4-dioxobutanoate

To a solution of diethyl oxalate (4.70 ml, 34.3 mmol) and 2-acetyl-4-methylpyridine (4.73 g, 34.3 mmol) in ethanol (49 mL) was added sodium ethoxide (21 wt.% in ethanol, 14.21 mL, 38.1 mmol). The reaction mixture was stirred at room temperature overnight. LC-MS indicated that the reaction went to completion. After the mixture was concentrated and diluted with water (50 mL), the aqueous layer was neutralized with acetic acid to pH = 5 and extracted with diethyl ether (50 mL × 3). The organic layers were washed with brine and dried over anhydrous sodium sulfate, filtered off, and concentrated in vacuo to yield ethyl 4-(4-methylpyridin-2-yl)-2,4-dioxobutanoate (7.66 g, 32.6 mmol, 95% yield). LC-MS (m/z): 236.1 $[M+H]^+$.

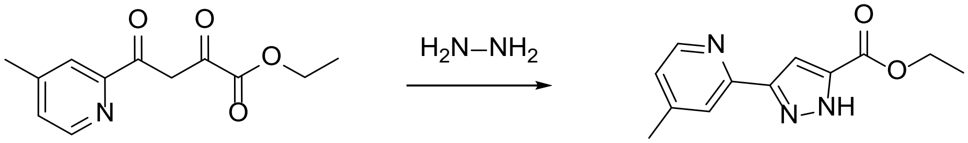

ethyl 5-(4-methylpyridin-2-yl)-1H-pyrazole-3-carboxylate

To a solution of ethyl 4-(4-methylpyridin-2-yl)-2,4-dioxobutanoate (6.66 g, 28.3 mmol) in ethanol (100 mL) was added hydrazine (0.933 mL, 29.7 mmol). The mixture was heated to 90 °C for 4 h. LC-MS indicated that the reaction didn't go to completion. After adding hydrazine (0.933 mL, 29.7 mmol), the mixture was heated for 1 h. After diluting with water (300 mL), the mixture was extracted with ether (300 mL × 2). The organic layers were dried over anhydrous sodium sulfate, filtered off, and concentrated in vacuo. The crude was purified by ISCO (0 to 10% methanol in

DCM) to yield ethyl 5-(4-methylpyridin-2-yl)-1H-pyrazole-3-carboxylate (5.1 g, 21.6 mmol, 76% yield). LC-MS (m/z): 232.2 $[M+H]^+$. $^1H$ NMR (500 MHz, DMSO-$d^6$) δ 14.13 (bs, 1H), 8.48 (d, $J$ = 5.0 Hz, 1H), 7.84 (s, 1H), 7.33 (s, 1H), 7.22 (d, $J$ = 4.7 Hz, 1H), 4.32 (q, $J$ = 7.1 Hz, 2H), 2.38 (s, 3H), 1.33 (t, $J$ = 7.1 Hz, 3H).

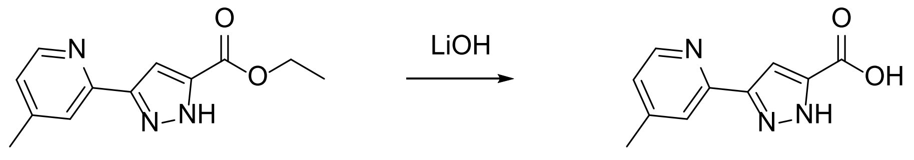

5-(4-methylpyridin-2-yl)-1H-pyrazole-3-carboxylic acid

To a solution of ethyl 5-(4-methylpyridin-2-yl)-1H-pyrazole-3-carboxylate (3.18 g, 13.75 mmol) in MeOH (10 mL) and water (10 mL) was added lithium hydroxide (0.692 g, 16.50 mmol). The mixture was heated at 65 °C for 1 h. After cooling down, pH of the mixture was adjusted to ~3 with 3 N HCl solution and extracted with EtOAc (50 mL × 2). The organic layers were dried over anhydrous sodium sulfate, filtered off, and concentrated in vacuo to yield 5-(4-methylpyridin-2-yl)-1H-pyrazole-3-carboxylic acid (2.6 g, 12.80 mmol, 93% yield). LC-MS (m/z): 204.2 $[M+H]^+$.

**Synthesis of final compounds**

**(*R*)-(3-benzylmorpholino)(3-(4-methylpyridin-2-yl)-1H-pyrazol-5-yl)methanone (compound 2)**

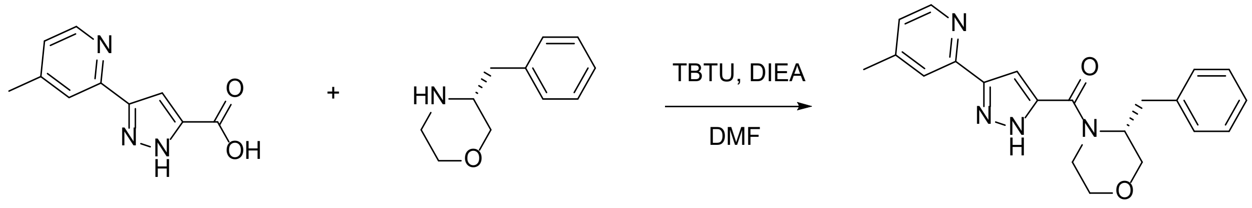

compound **2**

To a solution of 3-(4-methylpyridin-2-yl)-1H-pyrazole-5-carboxylic acid (20 mg, 0.098 mmol) and commercially available (*R*)-3-benzylmorpholine (20.93 mg, 0.118 mmol) in DMF (1 mL) was added DIEA (0.052 mL, 0.295 mmol) and TBTU (56.1 mg, 0.148 mmol) at 0 °C. The reaction mixture was stirred at room temperature for 2 h. LC-MS indicated that no starting materials remained. After adding water (3 mL), the reaction mixture was extracted with EtOAc (3 mL × 2). The combined organic layers were dried over anhydrous sodium sulfate, filtered off, and concentrated in vacuo. The crude product was purified by reverse phase HPLC under acidic conditions. The pure fractions were free-based with saturated sodium bicarbonate solution and extracted with EtOAc. The organic layers were dried over anhydrous sodium sulfate, filtered off, and concentrated in vacuo to yield (*R*)-(3-benzylmorpholino)(3-(4-methylpyridin-2-yl)-1H-pyrazol-5-yl)methanone (8 mg, 0.022 mmol, 22.2% yield). LC-MS (m/z): 363.3 $[M+H]^+$. HRMS (ESI) *m/z* calculated for C21H22N4O2 362.1743 $[M+H]^+$ found 363.1812. $^1H$ NMR (600 MHz, DMSO-$d^6$) δ 8.45 (d, $J$ = 5.0 Hz, 1H), 7.67 (s, 1H), 7.24 – 7.10 (m, 6H), 6.90 (s, 1H), 4.80 (s, 1H), 4.33 (s, 1H), 3.92 (d, $J$ = 7.9 Hz, 1H), 3.68 (d, $J$ = 11.6 Hz, 1H), 3.53 – 3.48 (m, 1H), 3.48 – 3.44

(m, 1H), 3.43 (s, 1H), 3.13 (dd, *J* = 13.5, 8.6 Hz, 1H), 3.02 (dd, *J* = 13.4, 6.6 Hz, 1H), 2.37 (s, 3H). $^{13}$C NMR (151 MHz, DMSO-*d*$^{6}$) δ 168.30, 148.45, 138.00, 137.74, 128.51, 128.25, 127.55, 127.49, 125.48, 125.40, 123.19, 71.55, 70.22, 66.85, 65.90, 48.82, 35.80, 34.44, 19.75.

**(*R*)-(3-(2-fluorobenzyl)morpholino)(3-(4-methylpyridin-2-yl)-1H-pyrazol-5-yl)methanone (compound 6)**

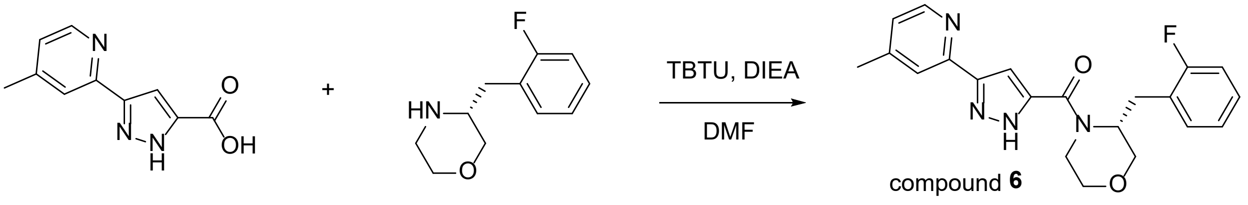

To a solution of 3-(4-methylpyridin-2-yl)-1H-pyrazole-5-carboxylic acid (260 mg, 1.28 mmol) and (R)-3-(2-fluorobenzyl)morpholine (275 mg, 1.41 mmol) in DMF (8 mL) was added DIEA (0.45 mL, 2.56 mmol) and TBTU (452 mg, 1.41 mmol) at 0 °C. The reaction mixture was stirred at room temperature for 16 h. LC-MS indicated that no starting materials remained. After adding water (30 mL), the reaction mixture was extracted with EtOAc (50 mL × 2). The combined organic layers were dried over anhydrous sodium sulfate, filtered off, and concentrated in vacuo. The crude product was purified by ISCO (0 to 10% MeOH in DCM) to yield (*R*)-(3-(2-fluorobenzyl)morpholino)(3-(4-methylpyridin-2-yl)-1H-pyrazol-5-yl)methanone (8 mg, 0.022 mmol, 22.2% yield). LC-MS (m/z): 381.2 $[M+H]^+$. HRMS (ESI) *m/z* calculated for C21H21FN4O2 380.1649 $[M+H]^+$ found 381.1719. $^{1}$H NMR (600 MHz, DMSO-*d*$^{6}$) δ 8.44 (d, *J* = 4.9 Hz, 1H), 7.65 (s, 1H), 7.27 (t, *J* = 7.6 Hz, 1H), 7.19 (q, *J* = 5.7 Hz, 1H), 7.14 (d, *J* = 5.0 Hz, 1H), 7.04 (t, *J* = 7.5 Hz, 1H), 7.00 (t, *J* = 9.3 Hz, 1H), 6.83 (s, 1H), 4.87 (s, 1H), 4.29 (d, *J* = 10.0 Hz, 1H), 3.92 (d, *J* = 7.4 Hz, 1H), 3.71 (d, *J* = 11.6 Hz, 1H), 3.58 (dd, *J* = 11.6, 3.2 Hz, 1H), 3.52 – 3.42 (m, 2H), 3.14 (dd, *J* = 13.8, 7.5 Hz, 1H), 3.08 (dd, *J* = 13.8, 7.7 Hz, 1H), 2.37 (s, 3H). $^{13}$C NMR (151 MHz, DMSO-*d*$^{6}$) δ 161.53, 161.16, 159.54, 148.44, 147.13, 131.04, 131.00, 127.68, 127.62, 124.47, 124.36, 123.47, 123.45, 123.07, 120.03, 114.35, 114.20, 104.54, 67.57, 65.94, 27.55, 19.75.

**(*R*)-(3-(2-chlorobenzyl)morpholino)(3-(4-methylpyridin-2-yl)-1H-pyrazol-5-yl)methanone (compound 7)**

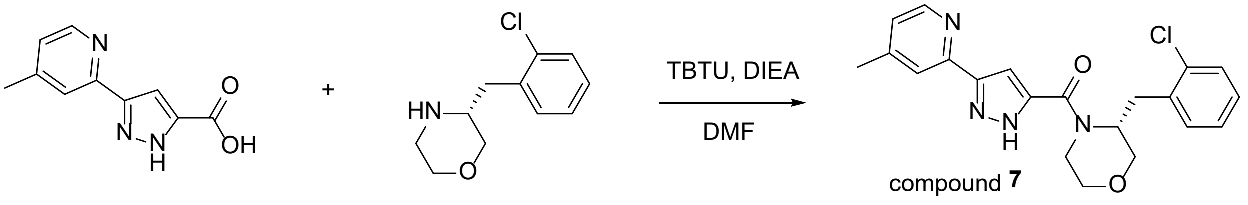

To a solution of 3-(4-methylpyridin-2-yl)-1H-pyrazole-5-carboxylic acid (30.0 mg, 0.148 mmol) and (*R*)-3-(2-chorobenzyl)morpholine (37.5 mg, 0.177 mmol) in DMF (3 mL) was added DIEA (0.077 mL, 0.443 mmol) and TBTU (52.1 mg, 1.62 mmol) at 0 °C. The reaction mixture was stirred at room temperature for 16 h. LC-MS indicated that no starting materials remained. After adding water (10 mL), the reaction mixture was extracted with EtOAc (20 mL × 2). The combined organic layers were dried over anhydrous sodium sulfate, filtered off, and concentrated in vacuo. The crude product was purified by preparative reverse phase HPLC to yield (*R*)-(3-(2-chlorobenzyl)morpholino)(3-(4-methylpyridin-2-yl)-1H-pyrazol-5-yl)methanone (45 mg, 0.086 mmol, 58.5% yield). LC-MS (m/z): 397.0 $[M+H]^+$. HRMS (ESI) *m/z* calculated for C21H21ClN4O2 396.1353 $[M+H]^+$ found 397.1413. $^1H$ NMR (600 MHz, DMSO-$d^6$) δ 8.48 (d, *J* = 5.1 Hz, 1H), 7.73 (s, 1H), 7.33 (dd, *J* = 7.0, 2.3 Hz, 1H), 7.29 (dd, *J* = 7.3, 2.0 Hz, 1H), 7.23 (dd, *J* = 5.2, 1.8 Hz, 1H), 7.22 – 7.15 (m, 2H), 6.87 (s, 1H), 4.92 (td, *J* = 7.5, 3.1 Hz, 1H), 4.31 – 4.26 (m, 1H), 3.99 – 3.93 (m, 1H), 3.75 (d, *J* = 11.6 Hz, 1H), 3.62 (dd, *J* = 11.7, 3.3 Hz, 1H), 3.59 – 3.46 (m, 2H), 3.26 (dd, *J* = 13.7, 7.2 Hz, 1H), 3.20 (dd, *J* = 13.7, 7.7 Hz, 1H), 2.42 (s, 3H). $^{13}C$ NMR (151 MHz, DMSO-$d^6$) δ 161.4, 148.3, 147.7, 147.6, 143.9, 143.7, 135.3, 133.1, 131.0, 128.5, 127.5, 126.3, 123.4, 120.4, 104.6, 67.7, 66.0, 51.0, 32.0, 19.9.

**(*R*)-(3-(2-methoxybenzyl)morpholino)(3-(4-methylpyridin-2-yl)-1H-pyrazol-5-yl)methanone (compound 8)**

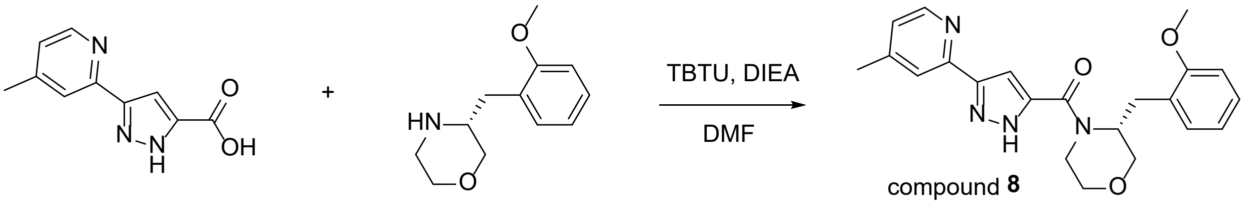

To a solution of 3-(4-methylpyridin-2-yl)-1H-pyrazole-5-carboxylic acid (30 mg, 0.148 mmol) and (R)-3-(2-methoxybenzyl)morpholine (30.6 mg, 0.148 mmol) in DMF (0.5 mL) was added DIEA (0.077 mL, 0.443 mmol) and TBTU (52.1 mg, 0.162 mmol) at 0 °C. The reaction mixture was stirred at room temperature for 15 h. LC-MS indicated that no starting materials remained. After adding water (2 mL), the reaction mixture was extracted with EtOAc (2 mL × 2). The combined organic layers were dried over anhydrous sodium sulfate, filtered off, and concentrated in vacuo. The crude product was purified by reverse phase HPLC under acidic conditions. The pure fractions were lyophilized to yield (*R*)-(3-(2-methoxybenzyl)morpholino)(3-(4-methylpyridin-2-yl)-1H-pyrazol-5-yl)methanone as its trifluoroacetic acid (TFA) salt (8.9 mg, 0.017 mmol, 11.3% yield). LC-MS (m/z): 393.3 $[M+H]^+$. HRMS (ESI) *m/z* calculated for C22H24N4O3 392.1848 $[M+H]^+$ found 393.1906. $^1H$ NMR (600 MHz, DMSO-$d^6$) δ 8.38 (s, 1H), 7.71 (s, 1H), 7.12 (q, *J* = 7.4 Hz, 2H), 7.03 (d, *J* = 5.0 Hz, 1H), 6.83 (d, *J* = 8.2 Hz, 1H), 6.82 – 6.75 (m, 2H), 4.94 (s, 1H), 4.44 (s, 1H), 3.89 (d, *J* = 7.9 Hz, 3H), 3.67 (s, 4H), 3.65 (d, *J* = 11.4 Hz, 3H), 3.52 (dd, *J* = 11.6, 3.1 Hz, 6H), 3.48 – 3.37 (m, 7H), 3.11 (dd, *J* = 13.4, 8.0 Hz, 3H), 2.96 (dd, *J* = 13.3, 7.1 Hz, 2H), 2.34 (s, 3H). Note: Broad peak between δH 2.7 – 4.2 ppm corresponds to the protons of residual water and results in inaccurate number of protons for the overlapped peaks.

**(*R*)-(3-(4-methylpyridin-2-yl)-1H-pyrazol-5-yl)(3-(pyridin-2-ylmethyl)morpholino)methanone (compound 5)**

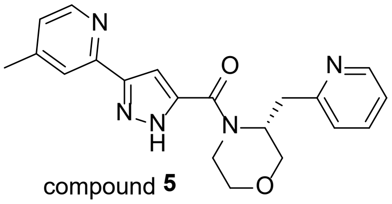

compound **5**

Compound **5** was prepared as its TFA salt by the above method using (*R*)-3-(pyridin-2-ylmethyl)morpholine. LC-MS (m/z): 364.2 [M+H]$^+$. HRMS (ESI) *m/z* calculated for C20H21N5O2 363.1695 [M+H]$^+$ found 364.1765. $^1$H NMR (600 MHz, DMSO-*d*$^6$) δ 8.51 (dd, *J* = 5.3, 1.8 Hz, 1H), 8.46 (d, *J* = 5.2 Hz, 1H), 7.88 (td, *J* = 7.6, 1.8 Hz, 1H), 7.73 (s, 1H), 7.48 (d, *J* = 7.9 Hz, 1H), 7.40 – 7.35 (m, 1H), 7.22 (d, *J* = 5.1 Hz, 1H), 6.94 (s, 1H), 5.01 (td, *J* = 7.4, 3.0 Hz, 1H), 4.32 – 4.26 (m, 1H), 3.96 – 3.90 (m, 1H), 3.82 (d, *J* = 11.6 Hz, 1H), 3.62 (dd, *J* = 11.6, 3.3 Hz, 1H), 3.58 – 3.45 (m, 2H), 3.40 – 3.30 (m, 2H), 2.40 (s, 3H). $^{13}$C NMR (151 MHz, DMSO-*d*$^6$) δ 161.2, 156.3, 148.6, 147.6, 147.5, 145.9, 143.7, 138.7, 124.4, 123.5, 121.9, 120.5, 104.8, 67.9, 65.9, 35.4, 19.9.

**(*S*)-(3-benzylmorpholino)(3-(4-methylpyridin-2-yl)-1H-pyrazol-5-yl)methanone (compound 12)**

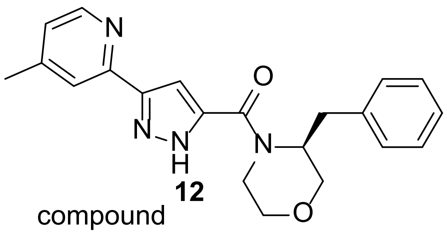

compound **12**

Compound **12** was prepared as its TFA salt by the above method using commercially available (*S*)-3-benzylmorpholine. LC-MS (m/z): 363.0 [M+H]$^+$. HRMS (ESI) *m/z* calculated for C21H22N4O2 362.1743 [M+H]$^+$ found 363.1814. $^1$H NMR (600 MHz, DMSO-*d*$^6$) δ 8.46 (d, *J* = 5.1 Hz, 1H), 7.72 (s, 1H), 7.25 – 7.18 (m, 5H), 7.18 – 7.12 (m, 1H), 6.93 (s, 1H), 4.79 – 4.73 (m, 1H), 4.30 – 4.24 (m, 1H), 3.95 – 3.90 (m, 1H), 3.68 (d, *J* = 11.6 Hz, 1H), 3.52 (dd, *J* = 11.6, 3.2 Hz, 1H), 3.50 – 3.40 (m, 2H), 3.13 (dd, *J* = 13.5, 8.6 Hz, 1H), 3.03 (dd, *J* = 13.5, 6.7 Hz, 1H), 2.40 (s, 3H). $^{13}$C NMR (151 MHz, DMSO-*d*$^6$) δ 161.3, 148.3, 147.8, 147.6, 137.7, 128.5, 127.6, 125.5, 123.4, 120.4, 104.7, 66.9, 65.9, 34.5, 19.9.

**(*R*)-(3-isopropylmorpholino)(3-(4-methylpyridin-2-yl)-1H-pyrazol-5-yl)methanone (compound 4)**

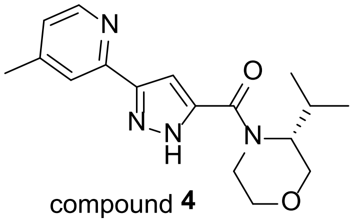

compound **4**

Compound **4** was prepared as its TFA salt by the above method using commercially available (*R*)-3-isopropylmorpholine. LC-MS (m/z): 315.3 [M+H]$^+$. HRMS (ESI) *m/z* calculated for C17H22N4O2 314.1743 [M+H]$^+$ found 315.1820. $^1$H NMR (600 MHz, DMSO-*d*$^6$) δ 8.45 (d, *J* = 5.0 Hz, 1H), 7.74 (s, 1H), 7.14 (d, *J* = 5.2 Hz, 1H), 7.04 (s, 1H), 4.33 (dd, *J* = 13.8, 3.1 Hz, 1H), 4.25 (d, *J* = 9.2 Hz, 1H), 3.98 (d, *J* = 11.8 Hz, 1H), 3.85 (dd, *J* = 11.3, 3.8 Hz, 1H), 3.52 (dd, *J* = 11.8, 3.2 Hz, 1H), 3.45 (td, *J* = 11.7, 3.1 Hz, 1H), 3.26 (s, 1H), 2.38 (s, 3H), 2.37 – 2.30 (m, 1H), 1.00 (d, *J* = 6.6 Hz, 3H), 0.89 (d, *J* = 6.8 Hz, 3H). $^{13}$C NMR (151 MHz, DMSO-*d*$^6$) δ 161.8, 148.5, 148.4, 147.0, 144.5, 122.9, 120.0, 104.7, 66.4, 66.0, 24.9, 19.7, 19.1, 18.2.

**(3-(4-Methylpyridin-2-yl)-1H-pyrazol-5-yl)(morpholino)methanone (compound 3)**

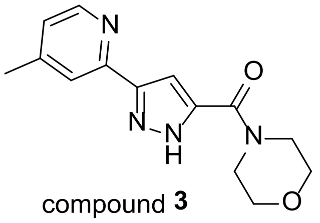

compound **3**

Compound **3** was prepared as its TFA salt by the above method using morpholine. LC-MS (m/z): 273.0 $[M+H]^+$. HRMS (ESI) *m/z* calculated for C14H16N4O2 272.1273 $[M+H]^+$ found 273.1346. $^1$H NMR (600 MHz, DMSO-$d^6$) δ 8.48 (d, *J* = 5.0 Hz, 1H), 7.81 (s, 1H), 7.22 (d, *J* = 5.0 Hz, 1H), 7.17 (s, 1H), 3.64 (s, 8H), 2.38 (s, 3H).

**(*R*)-(3-benzylmorpholino)(3-(pyridin-2-yl)-1H-pyrazol-5-yl)methanone (compound 14)**

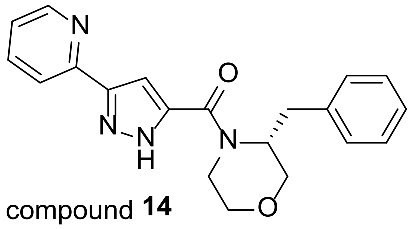

compound **14**

Compound **14** was prepared as its TFA salt by the above method using commercially available corresponding pyrazole acid and benzyl morpholine. LC-MS (m/z): 273.0 $[M+H]^+$. HRMS (ESI) *m/z* calculated for C20H20N4O2 348.1586 $[M+H]^+$ found 349.1651. $^1$H NMR (600 MHz, DMSO-$d^6$) δ 8.60 (dt, *J* = 4.8, 1.5 Hz, 1H), 7.86 – 7.80 (m, 2H), 7.30 (h, *J* = 4.3 Hz, 1H), 7.25 – 7.18 (m, 4H), 7.17 – 7.13 (m, 1H), 6.93 (s, 1H), 4.82 – 4.76 (m, 1H), 4.31 (d, *J* = 11.7 Hz, 1H), 3.95 – 3.90 (m, 1H), 3.68 (d, *J* = 11.6 Hz, 1H), 3.51 (dd, *J* = 11.6, 3.3 Hz, 1H), 3.50 – 3.39 (m, 2H), 3.13 (dd, *J* = 13.5, 8.6 Hz, 1H), 3.03 (dd, *J* = 13.5, 6.6 Hz, 1H). $^{13}$C NMR (151 MHz, DMSO-$d^6$) δ 161.5, 148.7, 148.3, 137.7, 136.3, 128.5, 127.6, 127.5, 125.5, 122.2, 119.4, 104.7, 66.9, 65.9, 34.5.

**(*R*)-(3-(2-fluoro-6-methoxybenzyl)morpholino)(3-(4-methylpyridin-2-yl)-1H-pyrazol-5-yl)methanone (compound 9) and (*S*)-(3-(2-fluorobenzyl)morpholino)(3-(4-methylpyridin-2-yl)-1H-pyrazol-5-yl)methanone (compound 10)**

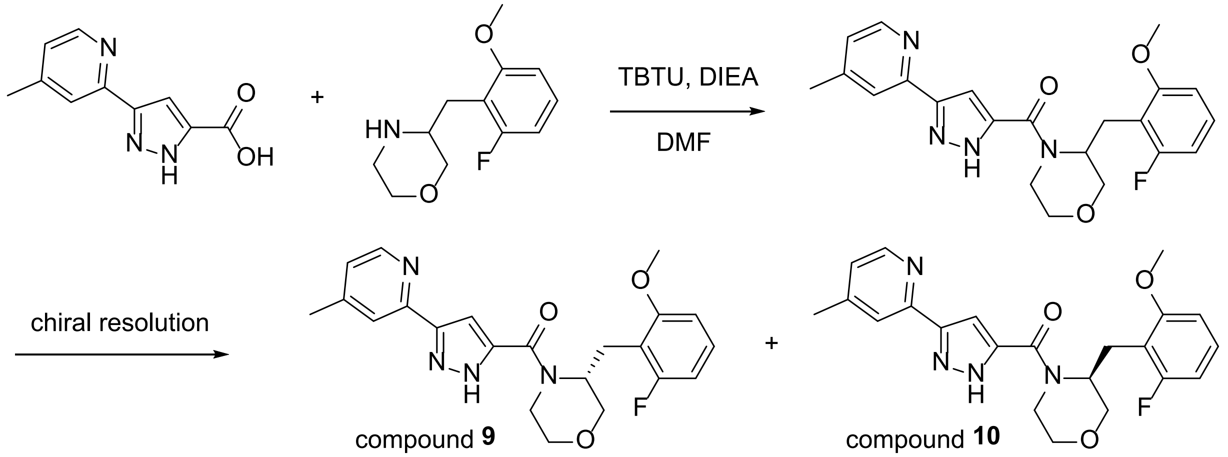

To a solution of 3-(4-methylpyridin-2-yl)-1H-pyrazole-5-carboxylic acid (80 mg, 0.394 mmol) and (+/-)-3-(2-fluoro-6-methoxybenzyl)morpholine (140 mg, 0.413 mmol) in DMF (2 mL) was added DIEA (0.206 mL, 1.18 mmol) and TBTU (152 mg, 0.472 mmol) at 0 °C. The reaction mixture was stirred at room temperature for 5 h. LC-MS indicated that no starting materials remained. After adding water (10 mL), the reaction mixture was extracted with EtOAc (20 mL × 2). The combined organic layers were dried over anhydrous sodium sulfate, filtered off, and concentrated in vacuo. The crude product was purified by reverse phase HPLC under acidic conditions. The pure fractions were free-based with saturated sodium bicarbonate solution and extracted with EtOAc. The organic layers were dried over anhydrous sodium sulfate, filtered off, and concentrated in vacuo to yield (+/-)-(3-(2-fluoro-6-methoxybenzyl)morpholino)(3-(4-methylpyridin-2-yl)-1H-pyrazol-5-yl)methanone (26 mg, 0.063 mmol, 16% yield). LC-MS (m/z): 411.1 $[M+H]^+$. HRMS (ESI) *m/z* calculated for C22H23FN4O3 410.1754 $[M+H]^+$ found 411.1814. $^1H$ NMR (600 MHz, DMSO-$d^6$) δ 8.43 (d, $J$ = 5.1 Hz, 1H), 7.69 (s, 1H), 7.18 – 7.09 (m, 2H), 6.70 (s, 1H), 6.67 (d, $J$ = 8.4 Hz, 1H), 6.62 (t, $J$ = 8.9 Hz, 1H), 4.95 (s, 1H), 4.31 (d, $J$ = 13.0 Hz, 1H), 3.92 (dd, $J$ = 11.0, 3.2 Hz, 1H), 3.75 (d, $J$ = 11.4 Hz, 1H), 3.69 (s, 3H), 3.64 (dd, $J$ = 11.4, 3.3 Hz, 1H), 3.56 – 3.42 (m, 3H), 2.54 (s, 2H), 2.38 (s, 3H). $^{13}C$ NMR (151 MHz, DMSO-$d^6$) δ 161.9, 161.8, 160.2, 158.6, 149.2, 148.4, 146.9, 144.7, 144.2, 127.4, 127.4, 122.6, 119.8, 113.3, 106.7, 106.5, 106.4, 104.3, 68.5, 66.0, 55.3, 40.4, 22.2, 19.8.

The racemic mixture of (+/-)-(3-(2-fluoro-6-methoxybenzyl)morpholino)(3-(4-methylpyridin-2-yl)-1H-pyrazol-5-yl)methanone (20 mg) was resolved by chiral supercritical fluid chromatography (SFC) to provide two enantiomers. The absolute configurations were determined on the basis of the biochemical $IC_{50}$ values and the X-ray co-crystallographic data. Chiral resolution conditions (SFC): AD column, SFC = 100 mL/min, $CO_2$/MeOH=70/30, 14 mg/2 mL MeOH, Load = 10 mg/1.5 mL, 236 bar.

**(*R*)-(3-(2-fluoro-6-methoxybenzyl)morpholino)(3-(4-methylpyridin-2-yl)-1H-pyrazol-5-yl)methanone (compound 9)** HRMS (ESI) *m/z* calculated for C22H23FN4O3 410.1754 $[M+H]^+$ found 411.1810. Retention time ($R_f$): 2.63 min (Peak 1), >99% *ee*; $IC_{50}$ = 2.5 μM

**(*S*)-(3-(2-fluoro-6-methoxybenzyl)morpholino)(3-(4-methylpyridin-2-yl)-1H-pyrazol-5-yl)methanone (compound 10)** HRMS (ESI) *m/z* calculated for C22H23FN4O3 410.1754 $[M+H]^+$ found 411.1801. Retention time ($R_f$): 4.68 min (Peak 2), 94.2% *ee*; $IC_{50}$ = 31.3 μM.

**(*R*)-(3-(2-chloro-6-methoxybenzyl)morpholino)(3-(4-methylpyridin-2-yl)-1H-pyrazol-5-yl)methanone (compound 11)**

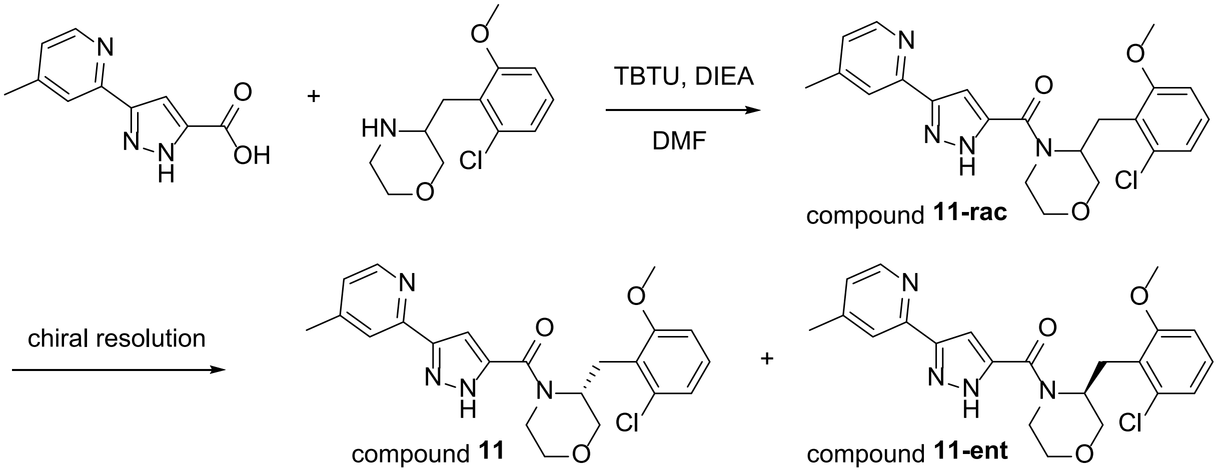

Compound **11-rac** was prepared by the above method using (+/-)-3-(2-chloro-6-methoxybenzyl)morpholine. Chiral resolution conditions (SFC): AD column, SFC = 100 mL/min, $CO_2$/isopropyl alcohol=70/30, 32 mg/4 mL MeOH, Load = 8 mg/1 mL, 266 bar

**(*R*)-(3-(2-chloro-6-methoxybenzyl)morpholino)(3-(4-methylpyridin-2-yl)-1H-pyrazol-5-yl)methanone (compound 11)** LC-MS (m/z): 427.3 $[M+H]^+$. HRMS (ESI) *m/z* calculated for C22H23ClN4O3 426.1459 $[M+H]^+$ found 427.1515. $^1$H NMR (600 MHz, DMSO-$d^6$) δ 8.43 (d, $J$ = 5.0 Hz, 1H), 7.67 (s, 1H), 7.14 – 7.08 (m, 2H), 6.86 (d, $J$ = 8.0 Hz, 1H), 6.81 (d, $J$ = 8.3 Hz, 1H), 6.67 (s, 1H), 5.03 – 4.97 (m, 1H), 4.34 – 4.29 (m, 1H), 3.94 (dd, $J$ = 11.2, 3.7 Hz, 1H), 3.77 (d, $J$ = 11.4 Hz, 1H), 3.72 (s, 3H), 3.65 (dd, $J$ = 11.4, 3.3 Hz, 1H), 3.63 – 3.55 (m, 1H), 3.46 (td, $J$ = 11.8, 3.0 Hz, 1H), 3.35 (dd, $J$ = 13.5, 8.7 Hz, 1H), 3.07 (dd, $J$ = 13.5, 5.6 Hz, 1H), 2.38 (s, 3H). $^{13}$C NMR (151 MHz, DMSO-$d^6$) δ 158.5, 148.3, 134.0, 127.5, 120.5, 109.2, 104.2, 68.8, 66.1, 55.3, 40.4, 26.8, 19.8. Retention time ($R_f$): 3.02 min (Peak 1), >99% *ee*; $IC_{50}$ (LpxA) = 0.46 μM

**(*S*)-(3-(2-chloro-6-methoxybenzyl)morpholino)(3-(4-methylpyridin-2-yl)-1H-pyrazol-5-yl)methanone (compound 11-ent)** LC-MS (m/z): 427.3 $[M+H]^+$. Retention time ($R_f$): 4.68 min (Peak 2), 96.7% *ee*; $IC_{50}$ (LpxA) = 1.39 μM.

**(*R*)-(3-(2-chloro-6-methoxybenzyl)morpholino)(3-(4-methylpyridin-2-yl)-1H-pyrazol-5-yl)methanone (compound 13)**

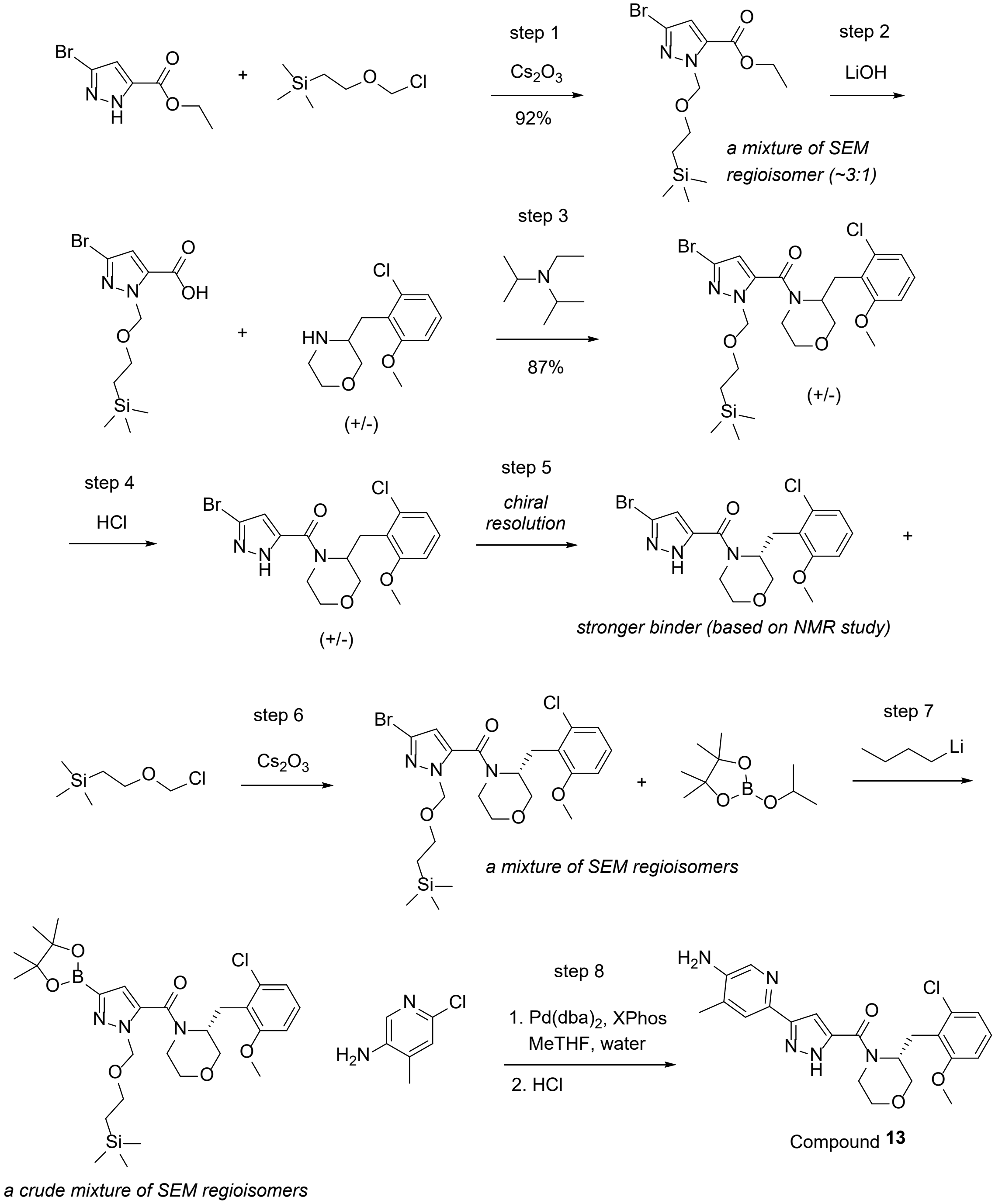

<u>Step 1</u>

**ethyl 3-bromo-1-((2-(trimethylsilyl)ethoxy)methyl)-1H-pyrazole-5-carboxylate and ethyl 5-bromo-1-((2-(trimethylsilyl)ethoxy)methyl)-1H-pyrazole-3-carboxylate**

To a solution of ethyl 3-bromo-1H-pyrazole-5-carboxylate (4.1 g, 18.72 mmol) in DMF (62.4 mL) was added cesium carbonate (9.15 g, 28.1 mmol). The reaction mixture was cooled down to 0 °C followed by addition of (2-(chloromethoxy)ethyl)trimethylsilane (3.65 mL, 20.59 mmol). The reaction mixture was warmed up to room temperature and stirred for 3 h. After quenched with water, the reaction mixture was extracted with EtOAc. The organic layer was dried over anhydrous sodium sulfate, filtered off, and concentrated in vacuo. The crude product was purified by ISCO (0% to 50% EtOAc in heptane) to yield a regioisomeric mixture of ethyl 3-bromo-1-((2-(trimethylsilyl)ethoxy)methyl)-1H-pyrazole-5-carboxylate and ethyl 5-bromo-1-((2-(trimethylsilyl)ethoxy)methyl)-1H-pyrazole-3-carboxylate (6 g, 17.18 mmol, 92% yield). LC-MS (m/z): 349, 351 $[M+H]^+$, 1.13 min, 1.2 min.

Step 2

**3-bromo-1-((2-(trimethylsilyl)ethoxy)methyl)-1H-pyrazole-5-carboxylic acid and 5-bromo-1-((2-(trimethylsilyl)ethoxy)methyl)-1H-pyrazole-3-carboxylic acid**

To a solution of a regioisomeric mixture of ethyl 3-bromo-1-((2-(trimethylsilyl)ethoxy)methyl)-1H-pyrazole-5-carboxylate and ethyl 5-bromo-1-((2-(trimethylsilyl)ethoxy)methyl)-1H-pyrazole-3-carboxylate (4.6 g, 13.17 mmol) in THF (26.3 mL) and MeOH (26.3 mL) was added 1 M LiOH aqueous solution (23.7 mL, 23.7 mmol) at room temperature. The reaction mixture was stirred at room temperature for 1 h. The pH of the reaction mixture was adjusted to ~4 by adding 1 N HCl solution. The reaction mixture was extracted with DCM (100 mL) and the organic layers were washed with water and brine, dried over sodium sulfate, filtered off, and concentrated in vacuo. The crude product was used in the next step without further purification. LC-MS (m/z): 321 $[M+H]^+$, 0.94 min and 0.99 min.

Step 3

**(+/-)-(3-bromo-1-((2-(trimethylsilyl)ethoxy)methyl)-1H-pyrazol-5-yl)(3-(2-chloro-6-methoxybenzyl)morpholino)-methanone and (+/-)-(5-bromo-1-((2-(trimethylsilyl)ethoxy)methyl)-1H-pyrazol-3-yl)(3-(2-chloro-6-methoxybenzyl)morpholino)methanone**

To a solution of 3-bromo-1-((2-(trimethylsilyl)ethoxy)methyl)-1H-pyrazole-5-carboxylic acid and 5-bromo-1-((2-(trimethylsilyl)ethoxy)methyl)-1H-pyrazole-3-carboxylic acid (2.05 g, 6.38 mmol), (+/-)-3-(2-chloro-6-methoxybenzyl)morpholine hydrochloride salt (2.008 g, 6.38 mmol), and DIEA (5.57 mL, 31.9 mmol) in DMF (21.27 mL) was added TBTU (3.07 g, 9.57 mmol) at 0 °C. The reaction mixture was stirred for 15 h at room temperature. After quenched with water, the reaction mixture was extracted with EtOAc. The organic layers were washed with water and brine, dried over anhydrous sodium sulfate, filtered off, and concentrated in vacuo. The crude product was purified by ISCO (gradient EtOAc in heptane) to yield a mixture of (+/-)-(3-bromo-1-((2-(trimethylsilyl)ethoxy)methyl)-1H-pyrazol-5-yl)(3-(2-chloro-6-methoxybenzyl)morpholino)-methanone and (+/-)-(5-bromo-1-((2-(trimethylsilyl)ethoxy)methyl)-1H-pyrazol-3-yl)(3-(2-chloro-6-methoxybenzyl)morpholino)methanone (3.03 g, 5.56 mmol, 87.2% yield). LC-MS (m/z): 544.3, 546.3 $[M+H]^+$, 1.19 min and 1.24 min.

Step 4

**(+/-)-(3-bromo-1H-pyrazol-5-yl)(3-(2-chloro-6-methoxybenzyl)morpholino)methanone**

To a solution of (+/-)-(5-bromo-1-((2-(trimethylsilyl)ethoxy)methyl)-1H-pyrazol-3-yl)(3-(2-chloro-6-methoxybenzyl)morpholino)methanone (3.03 g, 5.56 mmol) in DCM (1 mL) was added 4 M HCl in dioxane (10.51 mL, 42.0 mmol). The reaction mixture was stirred overnight. The volatile materials were removed in vacuo. The crude product was free-based with $NaHCO_3$ solution and extracted with EtOAc. The volatile materials were removed in vacuo to yield (+/-)-(3-bromo-1H-pyrazol-5-yl)(3-(2-chloro-6-methoxybenzyl)morpholino)methanone (1.67 g, 4.03 mmol, 72.5%). LC-MS (m/z): 414, 416 $[M+H]^+$, 0.74 min and 0.76 min.

Step 5

**(*R*)-(3-bromo-1H-pyrazol-5-yl)(3-(2-chloro-6-methoxybenzyl)morpholino)methanone**

The racemic mixture of (+/-)-(3-bromo-1H-pyrazol-5-yl)(3-(2-chloro-6-methoxybenzyl)morpholino)methanone (1.67 g) was resolved by chiral supercritical fluid chromatography (SFC) to provide (*R*)- or (*S*)-(3-bromo-1H-pyrazol-5-yl)(3-(2-chloro-6-methoxybenzyl)morpholino)methanone. The absolute configuration of two enantiomers was assigned on the basis of $^1H$-$^{13}C$ HMQC results of each enantiomer in the presence of LpxA product complex (the enantiomer from Peak 1 showed that NMR peaks are shifted as shown in the spectrum (right side) – the absolute stereochemistry of Peak 1 compound was assigned to *R*-configuration). Chiral resolution conditions (SFC): AD column, SFC = 100 mL/min, $CO_2$/MeOH=85/15, 278 mg/10 mL MeOH, Load = 27 mg/1.0 mL, 216 bar.

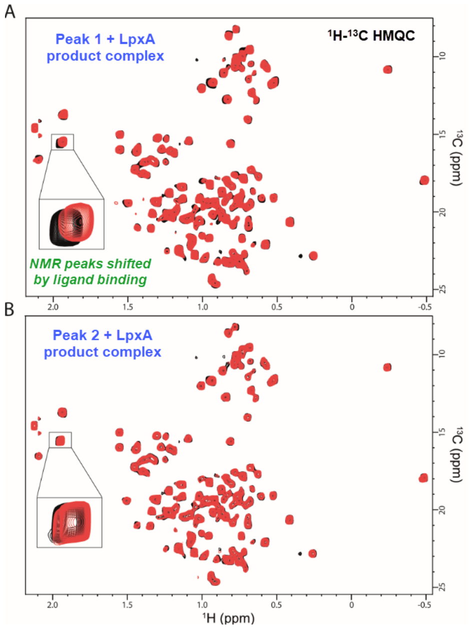

**(*R*)-(3-(2-fluoro-6-methoxybenzyl)morpholino)(3-(4-methylpyridin-2-yl)-1H-pyrazol-5-yl)methanone** Retention time ($R_f$): 5.01 min (Peak 1: 0.55 g, 32% yield), >99% *ee*, $^1H$ NMR (500 MHz, $CDCl_3$) δ 10.51 (s, 1H), 7.09 (t, J = 8.2 Hz, 1H), 6.88 (m, 1H), 6.61 (m, 1H), 6.25 (m, 1H), 4.59 (m, 1H), 4.28 – 3.43 (m, 7H), 4.13 (s, 3H), 2.98 (s, 1H).

**(*S*)-(3-(2-fluoro-6-methoxybenzyl)morpholino)(3-(4-methylpyridin-2-yl)-1H-pyrazol-5-yl)methanone** Retention time ($R_f$): 6.06 min (Peak 2), 87% *ee.*

Step 6

**(*R*)-(3-bromo-1-((2-(trimethylsilyl)ethoxy)methyl)-1H-pyrazol-5-yl)(3-(2-chloro-6-methoxybenzyl)morpholino)methanone and (*R*)-(5-bromo-1-((2-(trimethylsilyl)ethoxy)methyl)-1H-pyrazol-3-yl)(3-(2-chloro-6-methoxybenzyl)morpholino)methanone**

To a solution of (*R*)-(3-(2-fluoro-6-methoxybenzyl)morpholino)(3-(4-methylpyridin-2-yl)-1H-pyrazol-5-yl)methanone (0.55 g, 1.326 mmol) in DMF (4.42 mL) was added cesium carbonate

(0.648 g, 1.989 mmol). The reaction mixture was cooled down to 0 °C followed by addition of SEMCl (0.259 mL, 1.459 mmol). The reaction mixture was warmed up to room temperature and stirred for 3 h. After quenched with water, the reaction mixture was extracted with EtOAc. The organic layers were dried over anhydrous sodium sulfate, filtered off, and concentrated in vacuo. The crude product was purified by ISCO (0% to 50% EtOAc in heptane) to yield a regioisomeric mixture of (*R*)-(3-bromo-1-((2-(trimethylsilyl)ethoxy)methyl)-1H-pyrazol-5-yl)(3-(2-chloro-6-methoxybenzyl)morpholino)methanone and (*R*)-(5-bromo-1-((2-(trimethylsilyl)ethoxy)methyl)-1H-pyrazol-3-yl)(3-(2-chloro-6-methoxybenzyl)morpholino)methanone. LC-MS (m/z): 544, 546 $[M+H]^+$, 1.26 min and 1.30 min.

Step 7

**(*R*)-(3-(2-chloro-6-methoxybenzyl)morpholino)(3-(4,4,5,5-tetramethyl-1,3,2-dioxaborolan-2-yl)-1-((2-(trimethylsilyl)ethoxy)methyl)-1H-pyrazol-5-yl)methanone and (*R*)-(3-(2-chloro-6-methoxybenzyl)morpholino)(5-(4,4,5,5-tetramethyl-1,3,2-dioxaborolan-2-yl)-1-((2-(trimethylsilyl)ethoxy)methyl)-1H-pyrazol-3-yl)methanone**

To a solution of a mixture of (*R*)-(3-bromo-1-((2-(trimethylsilyl)ethoxy)methyl)-1H-pyrazol-5-yl)(3-(2-chloro-6-methoxybenzyl)morpholino)methanone and (*R*)-(5-bromo-1-((2-(trimethylsilyl)ethoxy)methyl)-1H-pyrazol-3-yl)(3-(2-chloro-6-methoxybenzyl)morpholino)methanone (532 mg, 0.976 mmol) and 2-isopropoxy-4,4,5,5-tetramethyl-1,3,2-dioxaborolane (239 µL, 1.172 mmol) in toluene (2.6 mL) and THF (651 µL) was slowly added 2.5 M n-butyllithium in hexanes (508 µl, 1.269 mmol) at -78 °C over 1.5 h through syringe pump. The reaction mixture was stirred at -78 °C for 2 h and allowed to warm up to room temperature. After stirring at room temperature for 1 h, the reaction mixture was quenched with saturated ammonium chloride solution and extracted with diethyl ether. The organic layers were dried over anhydrous sodium sulfate, filtered off, and concentrated in vacuo. The crude product was used in the next step without further purification. LC-MS (m/z): 510 $[M+H]^+$, 1.06 min (for boronic acid) and 593 $[M+H]^+$, 1.30 min (for boronic ester).

Step 8

**(*R*)-(3-(5-amino-4-methylpyridin-2-yl)-1H-pyrazol-5-yl)(3-(2-chloro-6-methoxybenzyl)morpholino)methanone**

To a solution of a mixture of (*R*)-(3-(2-chloro-6-methoxybenzyl)morpholino)(3-(4,4,5,5-tetramethyl-1,3,2-dioxaborolan-2-yl)-1-((2-(trimethylsilyl)ethoxy)methyl)-1H-pyrazol-5-yl)methanone and (*R*)-(3-(2-chloro-6-methoxybenzyl)morpholino)(5-(4,4,5,5-tetramethyl-1,3,2-dioxaborolan-2-yl)-1-((2-(trimethylsilyl)ethoxy)methyl)-1H-pyrazol-3-yl)methanone (264 mg, 0.446 mmol) in MeTHF (3.7 mL) and water (743 µl) was added $K_3PO_4$ (18.93 mg, 0.089 mmol), 1-(2-chloropyridin-4-yl)piperazine (140 mg, 0.982 mmol), XPhos (53.1 mg, 0.111 mmol), and $Pd(dba)_2$ (12.82 mg, 0.022 mmol). After purging with $N_2$ for 5 min, the reaction was heated at 110 °C for 1 h. After adding anhydrous sodium sulfate and EtOAc, the mixture was filtered off and the filtrate was concentrated in vacuo to yield crude coupling product as a regioisomeric mixture (LC-

MS (m/z): 581 $[M+H]^+$, 0.81 min). To the crude product in DCM (0.5 mL) was added 4 M HCl in dioxane (2.2 mL, 3.15 mmol). The reaction mixture was stirred for 16 h and the volatile materials were removed in vacuo. After adding sodium carbonate solution and EtOAc, the mixture was stirred for 15 min and extracted with EtOAc. The organic layers were washed with water and brine, dried over sodium sulfate, filtered off, and concentrated in vacuo. The crude product was purified by preparative SFC to yield (*R*)-(3-(5-amino-4-methylpyridin-2-yl)-1H-pyrazol-5-yl)(3-(2-chloro-6-methoxybenzyl)morpholino)methanone (8.4 mg, 0.017 mmol, 3.8% yield). LC-MS (m/z): 442.0 $[M+H]^+$. HRMS (ESI) *m/z* calculated for C22H24ClN5O3 441.1568 $[M+H]^+$ found 442.1633. $^1$H NMR (600 MHz, DMSO-$d^6$) δ 7.99 (s, 1H), 7.37 (s, 1H), 7.11 (t, *J* = 8.2 Hz, 1H), 6.86 (d, *J* = 8.0 Hz, 1H), 6.80 (d, *J* = 8.3 Hz, 1H), 6.43 (s, 1H), 5.01 (s, 1H), 4.27 (d, *J* = 13.7 Hz, 1H), 3.93 (dd, *J* = 11.3, 3.6 Hz, 1H), 3.76 (d, *J* = 11.4 Hz, 1H), 3.71 (s, 3H), 3.64 (dd, *J* = 11.5, 3.3 Hz, 1H), 3.57 (m, 1H), 3.45 (td, *J* = 11.8, 3.0 Hz, 1H), 3.33 (dd, *J* = 13.5, 8.7 Hz, 1H), 3.05 (dd, *J* = 13.5, 5.6 Hz, 1H), 2.17 (s, 3H). $^{13}$C NMR (151 MHz, DMSO-$d^6$) δ 158.5, 135.2, 134.0, 129.0, 127.5, 120.6, 109.2, 102.2, 68.9, 66.0, 55.3, 40.4, 39.9, 39.8, 39.7, 39.5, 39.4, 39.2, 39.1, 26.8, 15.7.

**Small molecule NMR spectroscopy**

$^1$H and $^{13}$C NMR spectra of compounds in DMSO-$d^6$ were acquired using Bruker 500 or 600 MHz AVANCE III HD systems equipped with QCI-F cryoprobe. Chemical shifts were calibrated using TMS as an internal standard. Elevated temperature of 393 K was adopted for compound **2** and compounds **4** – **13** to reduce severe line broadening observed at 300 K.

**High resolution LC-MS for compound validation**

ESI-MS data were recorded using a LTQ Qrbitrap mass spectrometer (Thermo Fisher Scientific) with electrospray ionization source. The resolution of the MS system was approximately 30000. The drug candidate was infused into the mass spectrometer by UPLC (Acquity, Waters) from sample probe. The separation was performed on Acquity UPLC BEH C18 1x50 mm column at 0.15 mL/min flow rate with the gradient from 5% to 95% in 2.8 min. Solvent A was water with 0.1% trifluoroacetic acid and solvent B was 75% methanol and 25% isopropyl alcohol with 0.1% trifluoroacetic acid. The mass accuracy of the system has been found to be <5 ppm.

**Variable-temperature NMR**

Variable-temperature 1D $^1$H NMR spectra of compound **6** in DMSO-$d^6$ were acquired using Bruker 400MHz AVANCE III HD system equipped with SmartProbe (Bruker Biospin, Billerica, MA, USA). Exchanging peak pairs were selected from isolated region of the spectra and analyzed for exchange rate using Dynamic NMR Line Shape Analysis module in TopSpin (Bruker Biospin, Billerica, MA, USA). The acquired exchange rates were plotted against temperature to extract activation energy on amide bond rotation by Arrhenius equation, which yielded $Ea = 16.6$ kcal/mol for compound **6**.

**Figure S1**

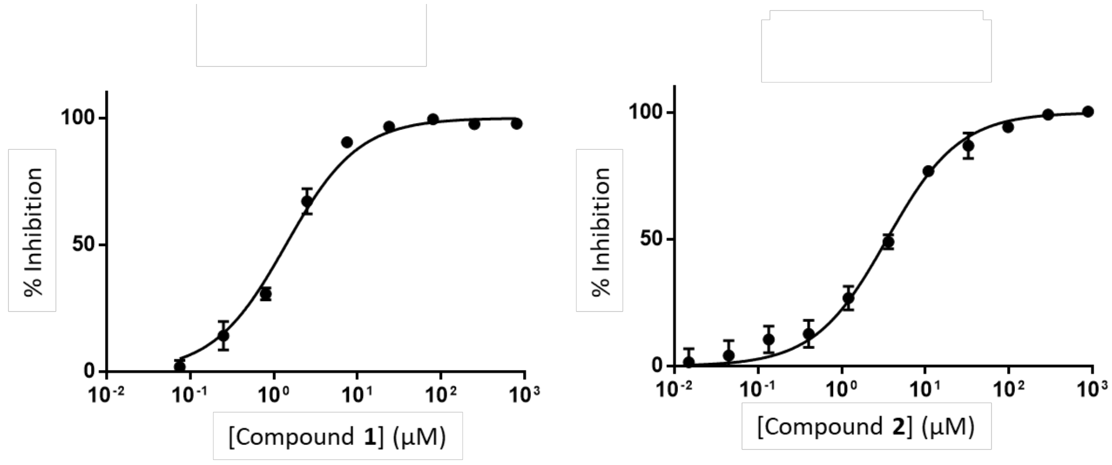

**Figure S1.** Biochemical $IC_{50}$ curves of compounds **1** and **2**.
Data are mean ± standard deviation of two assays for compound **1** and six representative assays for compound **2**.

**Figure S2**

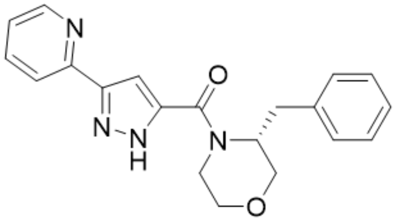

***Compound 14***
***(used for mutant selection)***

| | |
|---|---|
| MW | 348 |
| Biochemical $IC_{50}$ | 30 µM |
| *Ec* Δ*tolC* $IC_{50}$ | 12 µM |

**Figure S2.** Chemical structure and biological activities of compound **14**, the des-methyl analog of compound **2**, used for mutant selection.

**Figure S3**

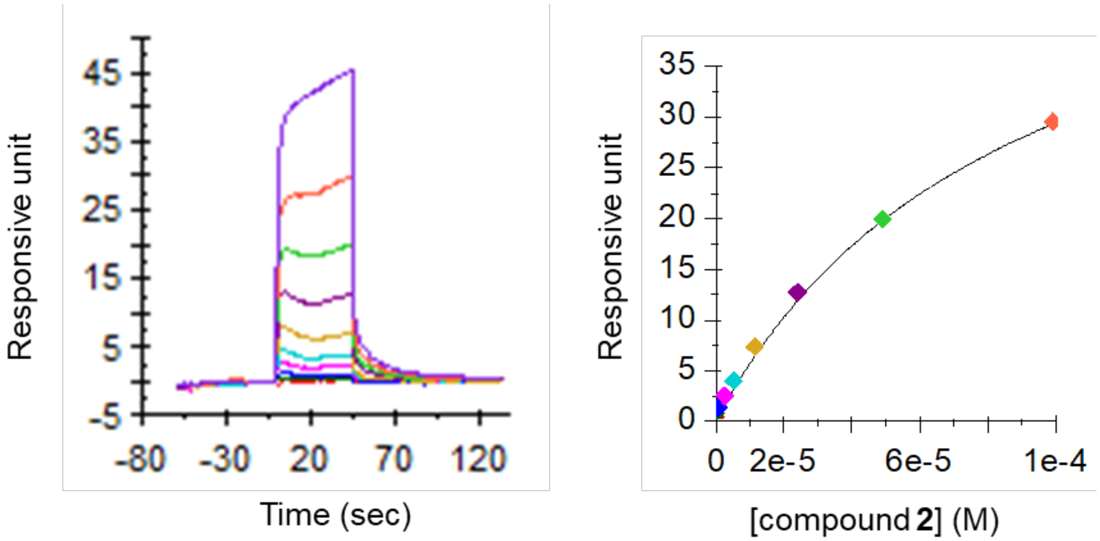

**Figure S3.** SPR sensogram (left) and dose response curve (right) for compound **2**.

**Figure S4**

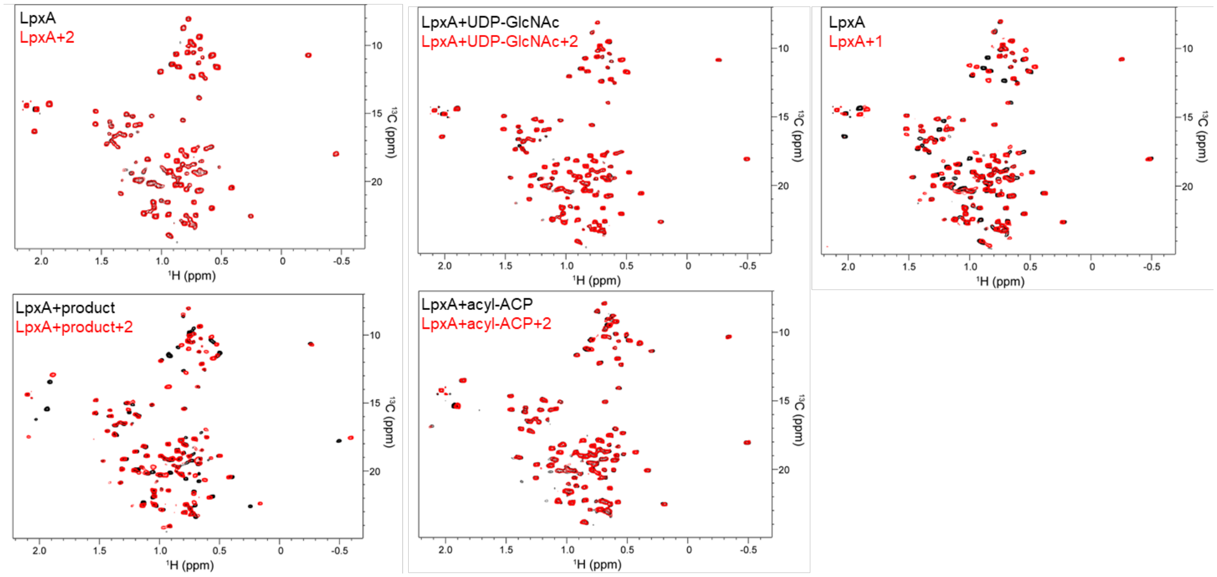

**Figure S4.** The methyl region of protein-observed 2D $^1$H-$^{13}$C HMQC spectra of the labeled amino acid residues (MILVAT) of LpxA**.** Each peak represents one methyl group of a Met, Ile, Leu, Val, Ala, or Thr residue LpxA. Figure 2 shows the zoomed images of the Met-methyl regions.

**Figure S5**

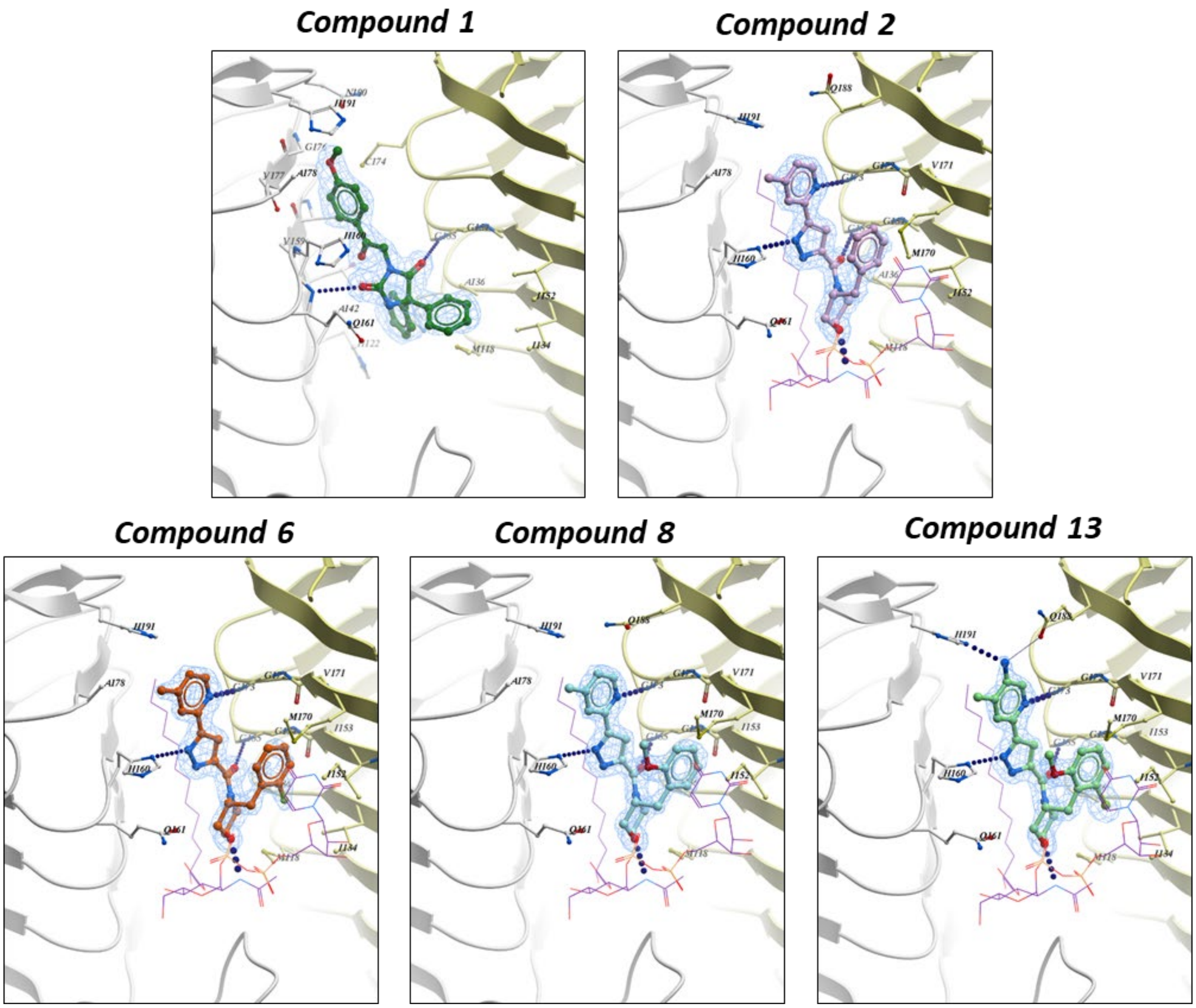

**Figure S5**. X-ray structures: compound **1**, **2**, **6**, **8** and **13**. A 2Fo-Fc density map (light blue, contoured at 3 sigma) for compound **2** bound to LpxA/LpxA product (1.8 Å resolution) is superimposed on the final refined model. LpxA is shown as ball and sticks with carbon colored white for protomer 1, orange for protomer 2, oxygen colored red and nitrogen colored blue. LpxA product is shown as lines with carbon colored purple. Compound **2** is shown as ball and sticks with carbon colored dark red.

**Table S1. Strains and plasmids used in this study**

| Strain/ plasmid | Description | Source/reference |
|---|---|---|
| *E. coli* strains | | |
| Top10 | *E. coli* cloning strain | Thermo Fisher |
| BL21-A1 | *$F^-$ ompT $hsdS_B$($r_B$-$m_B$-) gal dcm araB::T7RNAP-tetA($Tet^R$)* | Thermo Fisher |
| BL21 Star (DE3) | *$F^-$ ompT $hsdS_B$($r_B$-$m_B$-) gal dcm gal dcm rne-131 (DE3)* | Thermo Fisher |
| BL21(DE3) | *$F^-$ ompT $hsdS_B$($r_B$-$m_B$-) gal dcm (DE3)* | Thermo Fisher |
| BW25113 | *$F^-$ Δ(araD-araB)567 ΔlacZ4787::rrnB-3 $\lambda^-$ rph-1 Δ(rhaD-rhaB)568 hsdR514* | CGSC7926, Baba et al[21] |
| JW5503-1 | BW21553 *tolC*::$Km^r$ | CGSC11430, Baba et al[21] |
| MG1655 | *$F^-$ $\lambda^-$ ilvG- rfb-50 rph-1* | ATCC 47076 |
| VECO2526 | MG1655 Δ*tolC* | Lewis et al[22] |
| JW0451 | BW21553 *acrB*::$Km^r$ | CGSC8609, Baba et al [21] |
| imp4213 | *$lptD_{4213}$*($LptD_{\Delta 330\text{-}352}$) | Ruiz et al[23] |
| ARA0014 | *ΔtolC* FabZ(A146D) | This study |
| ARA0016 | *ΔtolC* FabZ(A78V) | This study |
| TUP0042 | *ΔtolC* FabZ(C139Y) | Ma et al[24] |
| TUP0074 | *ΔtolC* FabZ(P56L) | Ma et al[24] |
| TUP0115 | *ΔtolC* / pMMB206 (low copy, $Cm^r$, $P_{lac}$::*lacZα*) | Ma et al[24] |
| TUP0116 | *ΔtolC* / pTU457 (pMMB206 $P_{lac}$::*lpxA*) | Ma et al[24] |
| TUP0118 | *ΔtolC* / pTU507 (pMMB206 $P_{lac}$::*lpxC*) | Ma et al[24] |
| TUP0119 | *ΔtolC* / pTU433 (pMMB206 $P_{lac}$::*lpxD*) | Ma et al[24] |
| *E. coli* mobile plasmid collection | Plasmid clones of 4229 *E. coli* genes in JA200, $Ap^r$, IPTG-inducible, low copy, capable of plasmid transfer by F-mediated conjugation | National BioResource Project in Japan, Saka et al[4] |
| CDY0041 | BW25113 *ΔmdtBC ΔmdtF ΔmacB ΔentS ΔemrY ΔemrB ΔacrF ΔacrD* | Jones et al[25] |
| CDY0154 | BW25113 *ΔacrAB*::$Km^r$ *ΔmdtBC ΔmdtF ΔmacB ΔentS ΔemrY ΔemrB ΔacrF ΔacrD* | This study |
| TUP0092 | CDY0154 AccB(E128K) | This study |
| TUP0093 | CDY0154 LpxA(Q73L) | This study |
| TUP0097 | CDY0154 InfA(R66H) | This study |
| Plasmids | | |
| pKD46 | *R6K ori,* $Ap^r$, *frt*-$Km^r$-*frt* | Datsenko and Wanner[3] |
| pKD13 | *$repA101^{ts}$, $Ap^r$, Para::λRed genes (exo, bet, gam)* | Datsenko and Wanner[3] |
| pET-24a | *pBR $Km^r$ $lacI^q$ $P_{T7}$* | Novagen |
| pET-LpxA | *pBR $Km^r$ $lacI^q$ $P_{T7}$::lpxA-his6* | This study |
| pET-avi-LpxA | *pBR $Km^r$ $lacI^q$ $P_{T7}$::N-his6-3C-avi-lpxA* | This study |
| pET-AasS | *pBR $Km^r$ $lacI^q$ $P_{T7}$::Vh_aasS-his6* | Kreamer et al[5] |
| pTU448 | *pBR $Km^r$ $lacI^q$ $P_{T7}$::his6-tev-acp* | Kreamer et al[5] |
| pTU450 | *pBR $Km^r$ $lacI^q$ $P_{T7}$::his6-tev-acp_acpS* | Kreamer et al[5] |

### Table S2. Primers used in this study

| Primer # | Primer name | Sequence (5'-3') |
|---|---|---|
| E700 | Ec acrA pKD13 | ATTGACCAATTTGAAATCGGACACTCGAGGTTTACATATGATTCCGG GGATCCGTCGACC |
| E707 | acrB Rev-pkd13 | TTAGTGATTACACGTTGTATCAATGATGATCGACAGTATGTGTAGGC TGGAGCTGCTTCG |
| E466 | acrB +343-324 R | CCGCAGCAGGTAAAAGCAGT |
| E702 | Ec acrA -406-385 | ATACTGAGAGTGGATCGCCA |
| SP6 | Universal seq primer | GATTTAGGTGACACTATAG |
| TU117 | lpxDec 5'up | CTGTGAAATCCGTTGCCAACAG |
| TU243 | EclpxA HindIII | GGCCAGTGCCAAGCTTTTAACGAA TCAGACCGCGCGT |
| TU276 | EcLpxA -400F | ACTGGTGGATCGCGTGCTGG |
| TU332 | Ec FabZ US | CGGCCTGTCTCATTCTTACGATTGC |
| TU475 | EclpxA internal F | GTCAATTGGCGCGAACGCACACATTGG |
| TU475 | EclpxA internal F | GTCAATTGGCGCGAACGCACACATTGG |
| TU503 | infA F | GCCGGTTCAAATTACGGTAGTG |
| TU504 | infA R | CTCGTTCTTTCTCTTCGCCCATC |
| TU512 | accB F | CTGTCACAATCACACTAAAC |
| TU513 | accB R | TATCCAGCATGTTCGCCTCG |

### Table S3. X-ray data collection and refinement statistics

| | Compound 1 | Compound 2 | Compound 6 | Compound 8 | Compound 13 |
|---|---|---|---|---|---|
| RCSB ID | 6P9P | 6P9Q | 6P9R | 6P9S | 6P9T |
| Resolution range | 39.28 - 2.0 (2.072 - 2.0) | 32.38 - 1.7 (1.761 - 1.7) | 39.62 - 1.7 (1.761 - 1.7) | 25.81 - 1.75 (1.813 - 1.75) | 42.84 - 1.75 (1.813 - 1.75) |
| Space group | P 21 3 | P 21 3 | P 21 3 | P 21 3 | P 21 3 |
| Unit cell | 96.228 96.228 96.228 90 90 90 | 97.125 97.125 97.125 90 90 90 | 97.038 97.038 97.038 90 90 90 | 96.556 96.556 96.556 90 90 90 | 95.793 95.793 95.793 90 90 90 |
| Total reflections | 72508 (6808) | 178334 (2215) | 270242 (3224) | 263303 (6224) | 243346 (18334) |
| Unique reflections | 20319 (1985) | 31016 (1666) | 30050 (1392) | 30260 (2765) | 29813 (2956) |
| Multiplicity | 3.6 (3.4) | 5.7 (1.3) | 9.0 (2.3) | 8.7 (2.2) | 8.2 (6.2) |
| Completeness (%) | 99.48 (99.20) | 91.59 (50.20) | 88.94 (41.34) | 99.06 (91.71) | 99.92 (99.93) |
| Mean I/sigma(I) | 8.95 (1.10) | 20.34 (0.86) | 26.56 (1.92) | 19.54 (1.16) | 13.08 (0.89) |
| Wilson B-factor | 30.11 | 21.92 | 18.5 | 20.12 | 22.73 |
| R-merge | 0.1245 (1.196) | 0.05308 (0.5547) | 0.05591 (0.3703) | 0.07714 (0.6392) | 0.113 (1.588) |
| R-meas | 0.147 (1.421) | 0.05776 (0.7066) | 0.059 (0.4604) | 0.08174 (0.8076) | 0.1201 (1.736) |
| R-pim | 0.07662 (0.7576) | 0.02229 (0.4289) | 0.01844 (0.2683) | 0.02627 (0.4825) | 0.03982 (0.6933) |
| CC1/2 | 0.995 (0.346) | 0.997 (0.693) | 0.999 (0.787) | 0.999 (0.486) | 0.998 (0.509) |
| CC* | 0.999 (0.717) | 0.999 (0.905) | 1 (0.938) | 1 (0.809) | 0.999 (0.821) |
| Reflections used in refinement | 20245 (1982) | 30973 (1666) | 30034 (1392) | 30244 (2765) | 29800 (2955) |
| Reflections used for R-free | 1026 (95) | 1562 (61) | 1537 (80) | 1442 (136) | 1418 (144) |
| R-work | 0.1863 (0.2836) | 0.1643 (0.2866) | 0.1475 (0.2084) | 0.1545 (0.2848) | 0.1655 (0.2705) |
| R-free | 0.2232 (0.3120) | 0.1910 (0.3592) | 0.1770 (0.2277) | 0.1854 (0.2825) | 0.1850 (0.2925) |
| CC(work) | 0.970 (0.696) | 0.975 (0.735) | 0.972 (0.891) | 0.976 (0.733) | 0.972 (0.773) |
| CC(free) | 0.971 (0.702) | 0.964 (0.586) | 0.969 (0.869) | 0.962 (0.709) | 0.953 (0.759) |
| Number of non-hydrogen atoms | 2233 | 2445 | 2488 | 2473 | 2361 |
| macromolecules | 1987 | 2022 | 2022 | 2028 | 2022 |
| ligands | 52 | 104 | 106 | 105 | 108 |
| solvent | 194 | 319 | 360 | 340 | 231 |
| Protein residues | 263 | 263 | 263 | 263 | 263 |
| RMS(bonds) | 0.005 | 0.007 | 0.011 | 0.006 | 0.007 |
| RMS(angles) | 0.69 | 0.95 | 1.12 | 0.9 | 0.93 |
| Ramachandran favored (%) | 98.08 | 97.32 | 97.7 | 97.7 | 98.47 |
| Ramachandran allowed (%) | 1.92 | 2.68 | 2.3 | 2.3 | 1.53 |
| Ramachandran outliers (%) | 0 | 0 | 0 | 0 | 0 |
| Rotamer outliers (%) | 0 | 0 | 0 | 0 | 0 |
| Clashscore | 2.47 | 1.88 | 2.12 | 1.87 | 3.29 |
| Average B-factor | 34.25 | 26.62 | 21.86 | 23.2 | 26.54 |
| macromolecules | 32.91 | 24.25 | 19.35 | 20.73 | 24.87 |
| ligands | 55.65 | 33.19 | 24.94 | 25.25 | 28.56 |
| solvent | 42.18 | 39.54 | 35.04 | 37.26 | 40.17 |
| Number of TLS groups | 1 | 1 | 1 | 1 | 1 |

Statistics for the highest-resolution shell are shown in parentheses.
